# Type IV pili share a conserved mechanism of motor-independent retraction that is an inherent property of the pilus filament

**DOI:** 10.1101/2021.02.10.430637

**Authors:** Jennifer L. Chlebek, Lisa Craig, Ankur B. Dalia

## Abstract

Type IV pili (T4P) are dynamic surface appendages that promote virulence, biofilm formation, horizontal gene transfer, and motility in diverse bacterial species. Pilus dynamic activity is best characterized in T4P that use distinct ATPase motors for pilus extension and retraction. Many T4P systems, however, lack a dedicated retraction motor and the mechanism underlying this motor-independent retraction remains a mystery. Using the *Vibrio cholerae* competence pilus as a model system, we identify mutations in the major pilin gene that enhance motor-independent retraction. These mutants produced less stable pili, likely due to diminished pilin-pilin interactions within the filament. One mutation adds a bulky residue to α1C, a universally conserved feature of type IV pilins. We found that inserting a bulky residue into α1C of the retraction motor-dependent *Acinetobacter baylyi* competence T4P is sufficient to induce motor-independent retraction. Conversely, removing bulky residues from α1C of the retraction motor-independent *V. cholerae* toxin-co-regulated T4P stabilizes the filament and prevents retraction. Furthermore, alignment of pilins from the broader type IV filament (T4F) family indicated that retraction motor-independent T4P, Com pili, and type II secretion systems generally encode larger residues within α1C oriented toward the pilus core compared to retraction motor-dependent T4P. Together, our data demonstrate that motor-independent retraction relies on the inherent instability of the pilus filament that may be conserved in diverse T4Fs. This provides the first evidence for a long-standing, yet untested, model in which pili retract in the absence of a motor by spontaneous de-polymerization.

**SIGNIFICANCE:** Extracellular pilus filaments are critical for the virulence and persistence of many bacterial pathogens. A crucial property of these filaments is their ability to dynamically extend and retract from the bacterial surface. A detailed mechanistic understanding of pilus retraction, however, remains lacking in many systems. Here, we reveal that pilus retraction is an inherent property of the pilus filament. These observations are broadly relevant to diverse pilus systems, including those in many bacterial pathogens, and may help inform novel therapeutic strategies that aim to target pilus dynamic activity.

## INTRODUCTION

Type IV pili (T4P) are ubiquitous hair-like appendages that dynamically extend and retract from bacterial cells (1–5). These membrane-anchored nanomachines are important for the interactions of bacteria with their environment and facilitate activities like twitching motility, surface attachment, biofilm formation, protein secretion, interbacterial interactions, and horizontal gene transfer by natural transformation (2, 3, 6–11). The dynamic activity of T4P is central to many of these functions. T4P are also essential virulence factors in diverse pathogens (11–14). Thus, studying the regulation of these structures can uncover novel approaches for the development of therapeutic interventions.

The external pilus filament is a multimeric helical structure that is primarily composed of a single repeating subunit called the major pilin. This filament is built on an inner membrane platform protein commonly called PilC (15–17). Extension and retraction of the pilus is thought to occur through the interaction of cytoplasmic hexameric ATPases with the platform, whereby ATP hydrolysis facilitates coordinated conformational changes in the platform that promote polymerization or depolymerization of (18, 19).

There are three categories of T4P systems among Gram-negative bacteria: T4aP, T4bP, and T4cP. All T4P require an extension ATPase motor for pilus polymerization (20). A detailed mechanistic understanding of the factors that promote pilus retraction, however, remains lacking in many T4P. For T4aP, pilus depolymerization is mediated by dedicated retraction ATPase motors commonly referred to as PilT and PilU (21, 22). However, even in the absence of these retraction motors, numerous T4aP retain the ability to retract (4, 21, 23, 24). Many T4bP appear to lack a dedicated retraction ATPase, yet also retract (5, 25, 26), which suggests that diverse T4P may share a conserved mechanism for PilTU-independent retraction. Here, we investigated the mechanism of pilus retraction in the absence of dedicated retraction ATPases and find evidence to support a long-standing, yet untested, model in which pili retract via spontaneous depolymerization.

## RESULTS

### V. cholerae competence pili exhibit PilTU-independent retraction

To address the mechanism underlying PilTU-independent retraction, we employed the T4aP competence pili of *Vibrio cholerae* as a model system. These pili dynamically extend into the environment, bind DNA, and then retract to promote DNA uptake during horizontal gene transfer by natural transformation (1, 7). Ingested DNA can then be integrated into the genome by homologous recombination. To study the dynamic activity of *V. cholerae* competence pili, we can label pili using a technique in which an amino acid residue of the major pilin, PilA, is replaced with a cysteine (*pilA^S56C^*) for subsequent labeling with maleimide conjugated fluorescent dyes (2, 21, 27, 28). Using this technique, pili were directly observed by fluorescence microscopy. Functional dynamics of the competence pilus were also assessed through natural transformation assays.

The extension ATPase PilB and the retraction ATPases PilT and PilU drive the dynamic assembly and disassembly of *V. cholerae* competence pili, respectively. As expected, a Δ*pilB* mutant lacking the extension ATPase does not assemble pili (**Fig. 1A**) and is not transformable (**Fig. 1B**) (7, 21). As shown previously, a mutant strain lacking both retraction motors (Δ*pilTU*) is hyperpiliated and aggregates (8, 21) (**Fig. 1A**) yet it is still able to retract pili, albeit with lower retraction speed and frequency (21), which supports natural transformation at reduced levels (**Fig. 1A, B**). Importantly, previous work demonstrated that pilus retraction is required for DNA uptake during natural transformation for the Δ*pilTU* strain (1). Thus, the PilT and PilU motor ATPases are not absolutely required for pili to retract.

**Fig. 1.**
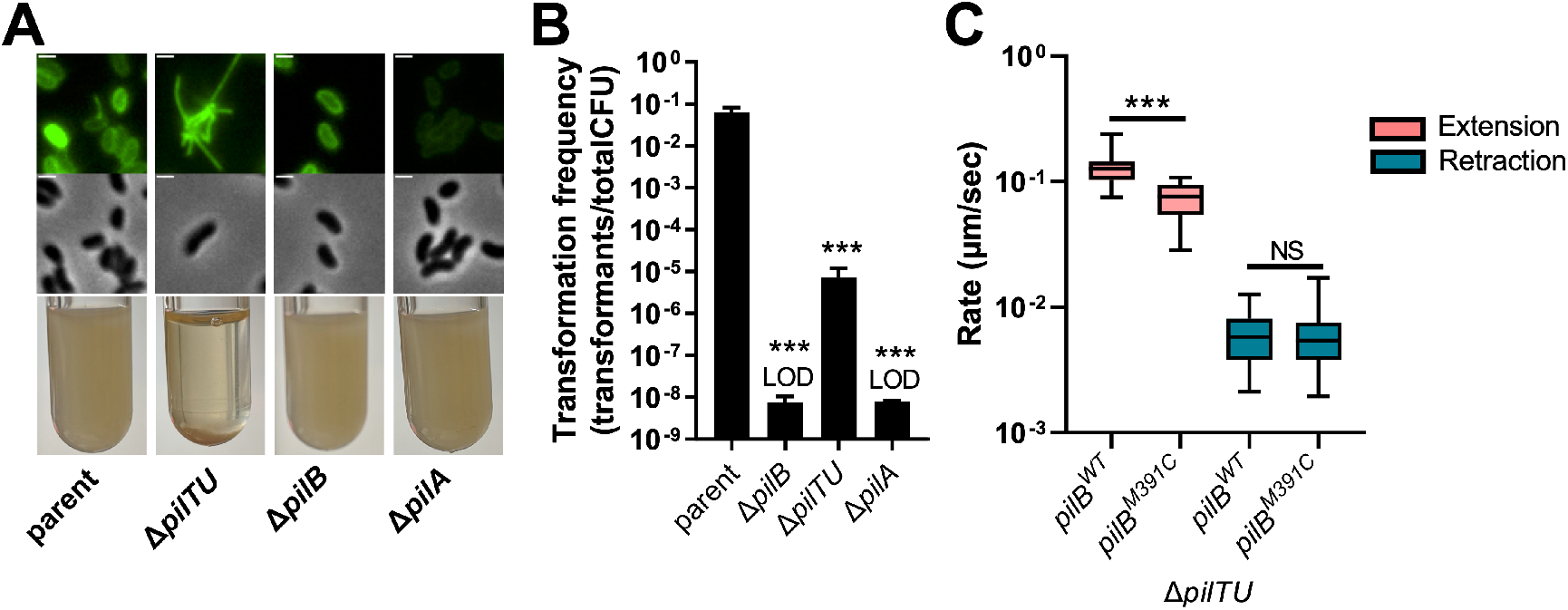
*V. cholerae* competence pili exhibit PilTU-independent retraction that is not powered by the extension motor PilB. **(A)** Representative images of surface piliation (top) and aggregation phenotypes (bottom) for the indicated strains. Scale bar, 1 μm. **(B)** Natural transformation assays of the indicated strains. Reactions were incubated with 200 ng of tDNA. All strains, *n* = 3. Data are shown as the mean ± SD. Asterisk(s) directly above bars denote comparisons to the parent strain. **(C)** Rates of extension (pink) and retraction (blue) for the indicated Δ*pilTU* strains was measured by epifluorescence time-lapse microscopy of AF488-mal labeled cells. To enhance dynamic activity, *pilB* alleles were placed under the control of a tightly inducible P_*BAD*_ promoter in a Δ*pilB* background and the expression of *pilB* was delayed until just before imaging was performed. Data are from three independent biological replicates. All strains, *n* = 44. Box plots represent the median and the upper and lower quartile, while the whiskers demarcate the range. All strains in **A-C** are derived from the parent strain, which contains a *pilA^S56C^* mutation to allow for pilus labeling. All comparisons were made by one-way ANOVA with Tukey’s post test. LOD, limit of detection; NS, not significant; *** = *P* < 0.001.

### PilB does not power PilTU-independent retraction

As mentioned above, many T4aP and T4bP are able to retract in the absence of dedicated retraction ATPases and the mechanism that mediates this PilTU-independent retraction remains unknown. T4cP also lack dedicated retraction motors, however these pilus systems use a bifunctional motor to power both pilus extension and retraction (29). Therefore, we sought to examine whether PilB powers PilTU-independent retraction of *V. cholerae* competence pili. Because PilB is required for pilus extension (**Fig. 1A**), we cannot test this using a Δ*pilB* mutant. Instead, we sought to determine if the ATPase activity of PilB was required for PilTU-independent retraction.

In order to test if PilB powers both extension and retraction via its ATPase activity, we employed the same technique described by Ellison *et al* to demonstrate that the ATPase CpaF powers both extension and retraction in *Caulobacter crescentus* T4cP. For that study, CpaF mutants that exhibit reduced ATPase activity slowed down the rates of both extension and retraction equally, indicating that CpaF likely powers both activities (29). Similarly, if *pilB* mutants that slow ATPase activity reduce the rates of both extension and PilTU-independent retraction, that would suggest that PilB is also a bifunctional ATPase. Measuring extension and retraction rates in a Δ*pilTU* strain, however, is exceedingly difficult because dynamic pilus events are rare in this hyperpiliated background (21). In order to increase the number of dynamic events, we generated a mutant strain that allowed us to tightly regulate the expression of PilB (Δ*pilTU* Δ*pilB* P_*BAD*_-*pilB*). By inducing PilB expression only during imaging, we greatly increase the number of pilus dynamic events observed (**Movies S1-3**).

To test if PilB is a bifunctional ATPase, we used a *pilB* mutant (*pilB^M391C^*) that was previously shown to reduce ATPase activity ~2-fold (30), and to correspondingly reduce the competence pilus extension rate ~2-fold (29). Consistent with these prior studies, Δ*pilTU* Δ*pilB* P_*BAD*_-*pilB^M391C^* exhibits ~2-fold slower extension rates compared to Δ*pilTU ΔpilB* P_*BAD*_-*pilB^WT^* (**Fig. 1C**) (29). However, retraction rates in Δ*pilTU* Δ*pilB* P_*BAD*_-*pilB^M391C^* were unaltered compared to Δ*pilTU* Δ*pilB* P_*BAD*_-*pilB^WT^* (**Fig. 1C**), which suggests that PilB is not a bifunctional motor. Importantly, previous work from our lab demonstrated that the ATPases required for polymerization of the other two T4P encoded by *V. cholerae* (MshE and TcpT) do not contribute to PilTU-independent retraction (21). Together, these results indicate that the retraction observed in the absence of PilTU is likely motor-independent. However, this does not reveal the mechanism underlying this motor-independent retraction.

### A forward genetic selection reveals a novel role for the major pilin, PilA, in motor-independent retraction

In order to identify the mechanism underlying motor-independent retraction we performed a forward genetic screen. As discussed above, mutants that exhibit enhanced retraction should also exhibit enhanced levels of natural transformation (**Fig. 1B**) (21). Thus, we reasoned that a suppressor screen selecting for mutants with enhanced levels of natural transformation might uncover gain-of-function mutations to components involved in motor-independent retraction. To that end, we recursively transformed a population of Δ*pilT* Δ*mutS* cells to enrich for mutants with enhanced rates of natural transformation. The deletion of *mutS* in this background increases the spontaneous mutation rate by disrupting the mismatch repair system (31). To maintain selection for each recursive round of natural transformation, different antibiotic resistance markers were swapped at a neutral locus. This selection was carried out in eleven biologically independent lineages. All lineages exhibited an enhanced transformation frequency following four rounds of selection. Single strains were isolated from each lineage and targeted sequencing of pilus genes was performed to identify potentially causative mutations. From the eleven populations, only 4 unique missense mutations were identified, all in the major pilin PilA (G34R, V74A, G78S, G80S). When these *pilA* mutations were introduced into a clean Δ*pilTU* genetic background, they all increased transformation frequency ~100-fold (**Fig. 2A**), confirming that they were sufficient to promote the enhanced rates of natural transformation observed in the original suppressor mutants.

**Fig. 2.**
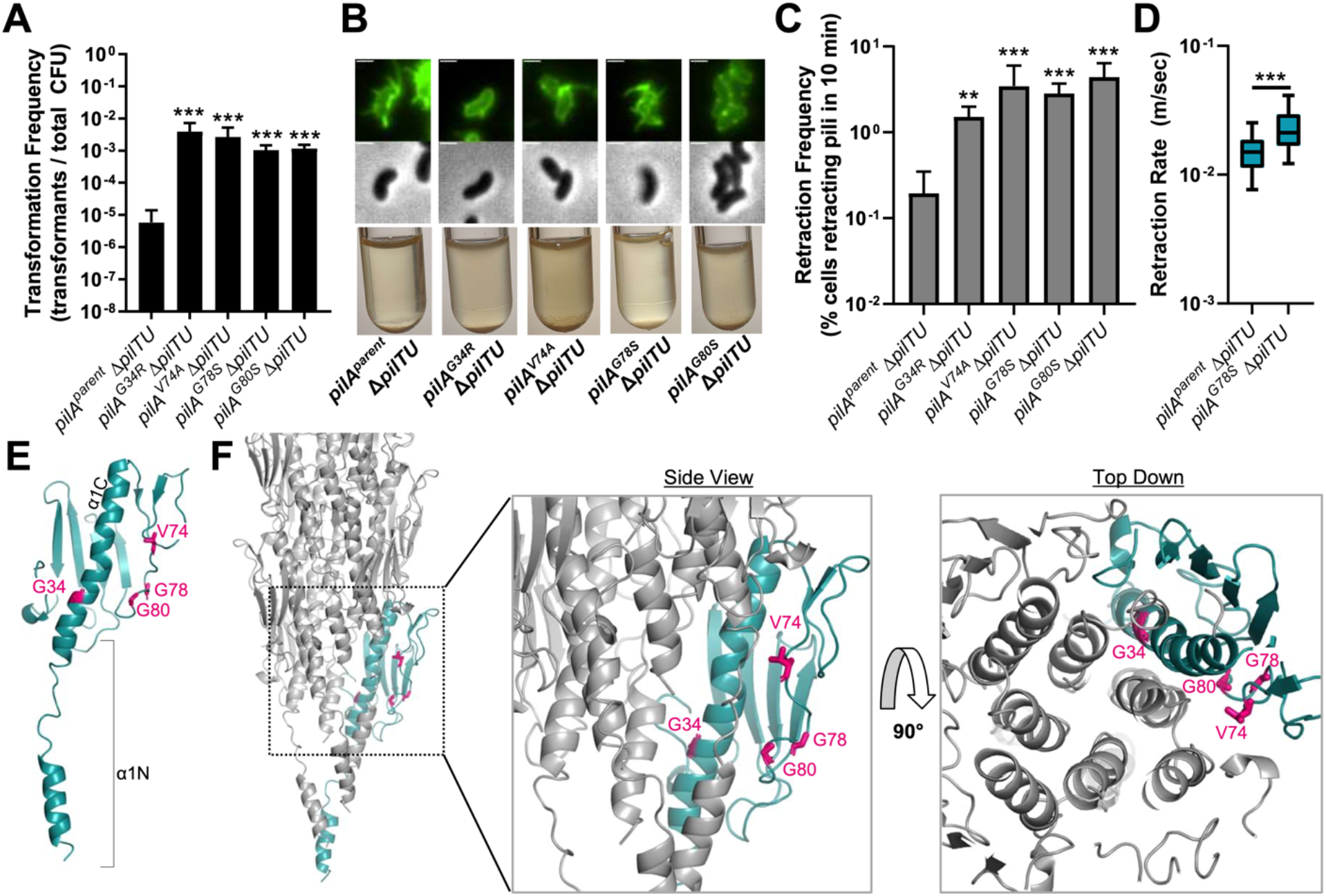
A forward genetic screen reveals mutations in PilA that enhance PilTU-independent pilus retraction. **(A)** Natural transformation assays of the indicated strains. Reactions were incubated with 200 ng of transforming DNA (tDNA). All strains, *n* = 3. **(B)** Representative images of surface piliation (top) and aggregation phenotypes (bottom) for the indicated strains. Scale bar, 1 μm. **(C)** Retraction frequency of the indicated strains measured by epifluorescence time-lapse microscopy of AF488-mal labeled cells. Data are presented as the percentage of cells within the population that exhibited a retraction event within the 10 min time-lapse. Data are from three independent biological replicates; *pilA^parent^* Δ*pilTU*, *n* = 1184. *pilA^G34R^* Δ*pilTU*, *n* = 718. *pilA^V74A^* Δ*pilTU*, *n* = 709. *pilA^G78S^* Δ*pilTU*, *n* = 864. *pilA^G80S^* Δ*pilTU*, *n* = 674. **(D)** Retraction rates of the indicated strains measured by epifluorescence time-lapse microscopy of AF488-mal labeled cells. Data are from three independent biological replicates; *pilA^parent^* Δ*pilTU, n* = 44. *pilA^G78S^* Δ*pilTU*, *n* = 30. **(E)** Structural model of PilA based on Phyre2 threading of the PilA sequence (NP_232053) onto PDB 1OQW (alignment coverage 80%, confidence 99.9%, identity 38%). Residues mutated in our suppressor screen are shown in stick representation and colored magenta. The α1N (residues 1-27) and α1C (residues 28-53) regions of PilA are also annotated. **(F)** Predicted model of the *V. cholerae* competence pilus in which the terminal pilin is shown in teal and other pilins in the fiber are shown in grey. The Phyre2-predicted PilA structure was superimposed onto one chain of the *Pseudomonas aeruginosa* PAK pilus (PDB 5VXY) and then the helical properties of the PAK pilus were used to generate a structural model for the PilA fiber. The PilA suppressor mutations are shown as magenta stick representations in the terminal pilin to highlight where they are within the fiber. All bar graphs are shown as the mean ± SD. All Box plots represent the median and the upper and lower quartile, while the whiskers demarcate the range. Asterisk(s) directly above bars denote comparisons to the parent strain. Comparisons in **A** and **C** were made by one-way ANOVA with Tukey’s post test. The comparison in **D** was made by Student’s t-test. ** = *P* < 0.01, *** = *P* < 0.001.

We also assessed the effect of these *pilA* suppressor mutations on natural transformation in a background where *pilTU* was intact. This analysis showed that all mutants transformed at levels similar to the parent strain (**Fig. S1A**), suggesting that these alterations to the major pilin did not impact pilus biogenesis or motor-dependent retraction. We directly imaged these pili using our pilus labeling approach. All *pilA* point mutants produced visible external filaments with and without *pilTU*, although three of the four strains with, *pilA^G34R^*, *pilA^V74A^* and *pilA^G80S^* mutations, produced shorter pili than the *pilA^parent^* strain (**Fig. 2B and Fig. S1B**). Even though the Δ*pilTU pilA^G34R^*, Δ*pilTU pilA^V74A^* and Δ*pilTU pilA^G80S^* strains produced shorter pili, all of these strains aggregate (**Fig. 2B**). Additionally, in a Δ*pilTU* background we used bulky maleimide conjugates (biotin-maleimide + neutravidin) to label pili to sterically obstruct pilus retraction (1, 2). This treatment prevented natural transformation, indicating that all of these *pilA* suppressor mutants still required pilus retraction to facilitate DNA uptake (**Fig. S2**). These data suggest that the *pilA* suppressor mutations enhance motor-independent retraction without altering pilus biogenesis or function.

The increased transformation frequencies observed for the suppressor mutations in a Δ*pilTU* background could result from increased retraction events or from increased rates of retraction. To distinguish between these possibilities, we directly observed pilus dynamic activity in strains with labeled pili using time-lapse epifluorescence microscopy. The retraction frequency could be assessed in all mutant backgrounds, however the rate of pilus retraction could only be accurately measured in the *pilA^parent^* and *pilA^G78S^* strains due to the short length of pilus filaments in the *pilA^G34R^*, *pilA^V74A^* and *pilA^G80S^* strains. These data revealed that pilus retraction frequency increased ~10-fold for all of the Δ*pilTU pilA* suppressor mutants, and that Δ*pilTU pilAG78S* pili retract ~2-times faster compared to the Δ*pilTU pilA^parent^* strain (**Fig. 2C, D**). We previously found that altering retraction rates even 7-fold has no discernable effect on transformation (21). Thus, the observed increase in the motor-independent retraction frequency likely explains the increase in transformation rates seen in these *pilA* suppressor mutants. The discrepancy between the 10-fold increase in retraction frequency and the 100-fold increase in transformation frequency may be attributed to the fact that many short dynamic pilus events occur that are not resolved by our microscopy-based approach.

Interestingly, in a background where the *pilTU* genes are intact, *pilA^G78S^* has no discernable effect on pilus retraction rates (**Fig. S1C**). Thus, our data indicate that the *pilA* suppressor mutations only affect pilus dynamics in the Δ*pilTU* background and did not affect pilus dynamics when *pilTU* were intact (**Fig. 2A, C** and **Fig. S1A, C**). This suggests that these *pilA* mutations only affect motor-independent pilus retraction and that their effect on pilus dynamics can be overcome by retraction motors.

### The stability of the pilus filament is a critical determinant for motor-independent retraction

In order to understand how these amino acid changes in PilA might increase the frequency of motor-independent retraction, we sought to understand how they might affect the overall structure of the major pilin and the pilus filament. To that end, we generated a model for PilA and for the competence pilus filament based on existing homologous structures. The predicted PilA structure was generated based on the *Pseudomonas aeruginosa* major pilin structure (Protein Data Bank (PDB) ID 1OQW) using Phyre2 (32, 33). The competence pilus model was generated by superimposing the PilA model onto one of the subunits in the *P. aeruginosa* pilus structure (PDB 5VXY) and then imposing helical symmetry to generate the multi-subunit filament (34). Additionally, the N-terminal half of the extended α-helix, α1N, of PilA was replaced with that of the *P. aeruginosa* pilin to reproduce the partial melting this α-helix undergoes when incorporated into the pilus (34, 35). The model of *V. cholerae* PilA shows the residue positions of the suppressor mutations: Gly34 is located in α1C, the C-terminal half of the primary N-terminal α-helix (α1); the other positions Val74, Gly78 and Gly80 are located in the αβ-loop that connects α1 with the central β-sheet of the globular domain (**Fig. 2E**). The pilus filament model revealed that all four suppressor mutation positions that enhance motor-independent retraction (PilA^G34R^, PilA^V74A^, PilA^G78S^, and PilA^G80S^) lie at interfaces between pilin subunits (**Fig. 2F).** When pilins are assembled into a filament, a portion of the α1 of each pilin is melted within a conserved region between residues 14-22, presumably to allow α1N to pack within the filament (34, 35). All of the PilA suppressor mutations encode residues that potentially contact α1N of a neighboring pilin subunit. Thus, altering contacts in this area might disrupt subunit packing and destabilize the pilus. We note that three of the four suppressor mutations in the *pilA* gene encode substitutions at glycines. Glycine has the smallest side chain of all amino acids, which allows greater backbone flexibility and is thus known as a “helix-breaker”. While Val74 is replaced with a smaller alanine, Gly34 is replaced by a bulky positively charged arginine, and Gly78 and Gly80 are both replaced with polar serines. The increase in bulk, polarity and/or the decrease in flexibility at these sites may disrupt both the backbone conformation at α1C and the αβ-loop as well as the close packing of the subunits, destabilizing the pilus filament.

How might alterations in pilin subunit interactions increase motor-independent retraction in these *pilA* suppressor mutants? It has previously been suggested that pili can spontaneously depolymerize in the absence of motors (5, 36–38), however, conclusive evidence for this model has been elusive. We hypothesize that a pilus with weaker pilin-pilin interactions would allow newly added subunits at the base of the pilus to diffuse more readily back into the membrane if pilus assembly is stalled, thereby facilitating motor-independent retraction through spontaneous depolymerization (see **Discussion** for a detailed description of this model). A consequence of weaker pilin-pilin interactions may be altered stability of the pilus filament. To test whether or not the suppressor mutants in fact produce less stable pili, we purified labeled pili and tested their thermal and chemical stability as well as their protease susceptibility, as previously described (38). Though we were unable to purify the very short PilA^G34R^, PilA^V74A^, and PilA^G80S^ filaments in sufficient quantities for these assays, we obtained testable amounts of PilA^parent^ and PilA^G78S^ pili from a Δ*pilTU* background. We found that the PilA^G78S^ filaments have similar chemical stability to the PilA^parent^ filaments (**Fig. S3**) but are less thermally stable and more susceptible to proteases (**Fig. 3A**). This instability may facilitate spontaneous depolymerization, thereby increasing the frequency of motor-independent retraction.

**Fig. 3.**
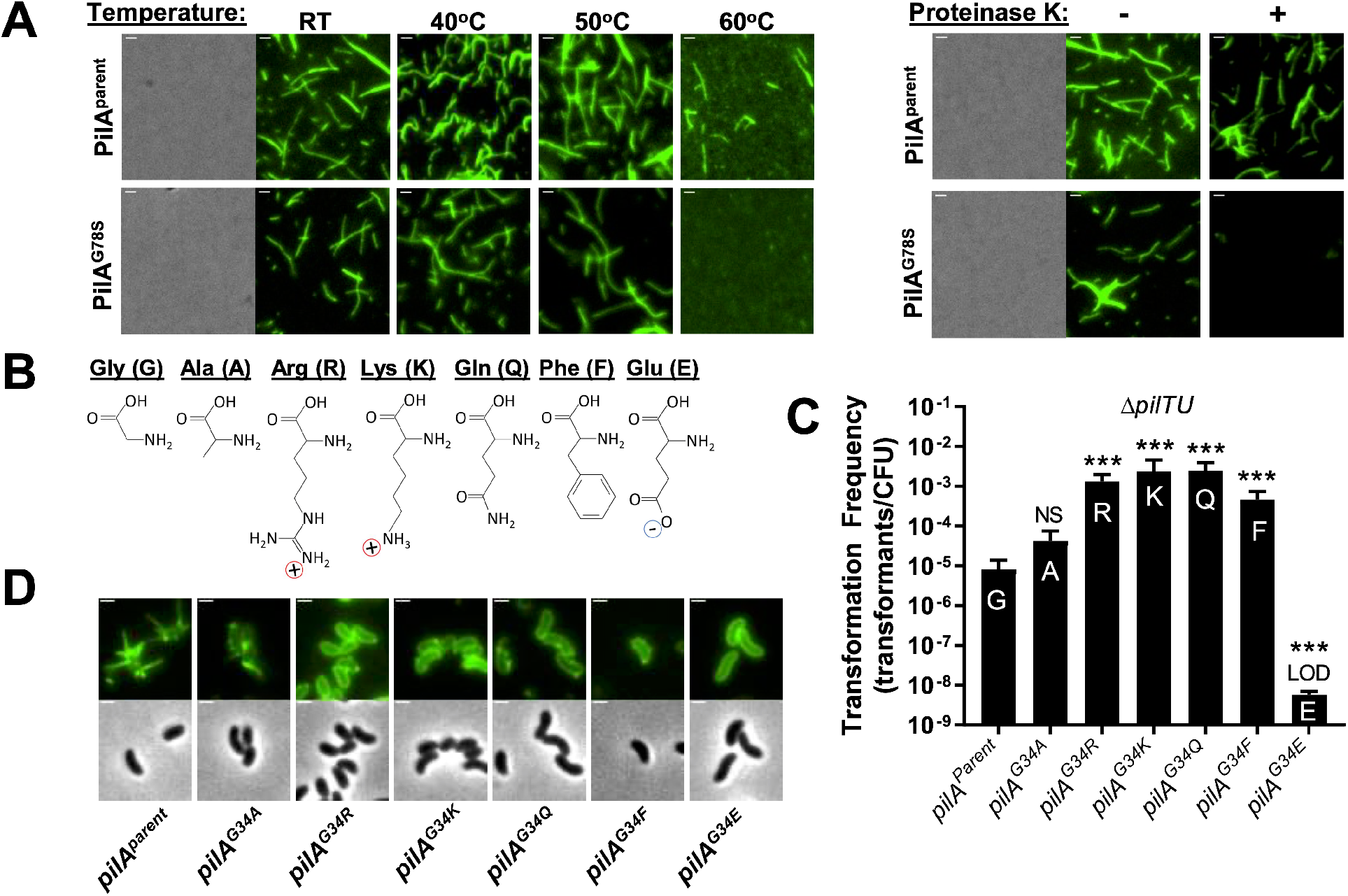
The stability of the pilus filament and the size of residues in α1C influence motor-independent retraction. **(A)** Stability assay comparing thermal stability and protease susceptibility of purified AF488-mal labeled pilus filaments from Δ*pilTU* strains expressing the indicated *pilA* allele. A representative phase-contrast image is shown to highlight that pilus preps are cell free. Fluorescent images show the presence or absence of intact pili under the indicated conditions. All images are representative of three independent biological replicates. Scale bar, 1 μm. **(B)** Skeletal formulas of the amino acids substituted at PilA^G34^ to highlight their chemical properties. **(C)** Natural transformation assays of the indicated Δ*pilTU* strains. Reactions were incubated with 200 ng of tDNA. For ease of mutant generation, the native *pilA* gene was knocked out and the indicated *pilA* allele was expressed at an ectopic location (under the control of an IPTG-inducible P_*tac*_ promoter). Data are shown as the mean ± SD and comparisons were made by one-way ANOVA with Tukey’s post test. Asterisk(s) directly above bars denote comparisons to the parent strain. All strains, *n* = 3. LOD, limit of detection; NS, not significant; *** = *P* < 0.001. **(D)** Representative images of surface piliation for the Δ*pilTU* strains used in **C**. Scale bar, 1 μm.

### Insertion of bulky residues in α1C of the V. cholerae competence pilus enhances motor-independent retraction

Since these data suggest that altering pilin-pilin interactions reduces filament stability and enhances motor-independent pilus retraction, we sought to understand the molecular mechanism of filament destabilization. We focused on the PilA^G34R^ mutation as it is located in a region of the major pilin, α1C, that is conserved among all major pilins from the type IV filament (T4F) family (**Fig. S4**) (20). The PilA^G34R^ mutation changes a small non-polar amino acid (glycine) to a large, positively charged amino acid (arginine). To determine whether the size or charge of the arginine side chain is responsible for filament destabilization and enhanced motor-independent retraction, we made a panel of mutants with amino acid substitutions at the Gly34 position in PilA (**Fig. 3B**). We expected that if the positive charge of the arginine was the critical feature, substitution with lysine but not other amino acids should enhance motor-independent retraction. However, if the size of the arginine is the important feature, substitution with other large amino acids, like glutamine and phenylalanine, should also phenocopy the *pilA^G34R^* mutant. Transformation assays showed that Δ*pilTU pilA^G34K^*, Δ*pilTU pilA^G34Q^* and Δ*pilTU pilA^G34F^* all phenocopied Δ*pilTU pilA^G34R^*, indicating that it is the size of the amino acid at position 34 and not its charge that is critical for motor independent retraction (**Fig. 3C**). Since substitutions with glutamine (polar) and phenylalanine (non-polar) both enhanced transformation similarly, this suggests that residue polarity at the Gly34 position in PilA is also not a critical feature for enhancing motor-independent retraction. Each of these mutants also exhibited a reduced pilus length compared to Δ*pilTU pilA^parent^*, similar to the original Δ*pilTU pilA^G34R^* mutant (**Fig. 3D**). Changing the Gly34 position to a negatively charged residue (*pilA^G34E^*) abolished transformation (**Fig. 3C**), most likely due to disruption of pilus biogenesis, which is supported by a lack of observable pilus filaments in this background (**Fig. 3D**). Substituting glycine with another small amino acid, alanine (*pilA^G34A^*), which is common in α-helices, only had a minor effect on transformation and a minimal effect on pilus length (**Fig. 3C, D**). Together, these data indicate that bulky residues at position 34 in PilA destabilize the pilus filament to enhance the frequency of spontaneous motor-independent retraction events.

### Residue bulkiness in α1C is sufficient to induce motor-independent retraction of the A. baylyi competence pilus

We next sought to determine whether the bulkiness at position 34 correlates with the ability of other T4P to retract spontaneously. First, we tested whether bulkiness in α1C was sufficient to promote motor-independent retraction in the naturally retraction motor-dependent competence T4aP of *Acinetobacter baylyi.* Like *V. cholerae* PilA, the *A. baylyi* major pilin ComP has a small amino acid, a serine, at position 34 (**Fig. S4**). *A. baylyi* competence pili can be studied using the same cysteine knock-in (*comP^T123C^*) and maleimide dye labeling approach described above (39), and we can functionally assess competence pilus dynamic activity through natural transformation assays. Unlike *V. cholerae*, *A. baylyi* Δ*pilTU* mutants do not exhibit any residual natural transformation (**Fig. 4A**) (21), which is consistent with an absence of motor-independent retraction for this T4aP. We hypothesized that introducing a bulky residue at position 34 of ComP would disrupt pilin-pilin interactions, destabilizing the pilus to promote motor-independent retraction. To that end, we substituted Ser34 of the *A. baylyi* major pilin ComP with arginine (*comP^S34R^*). The *comP^S34R^* strain produces pilus filaments, indicating that pilus biogenesis is not inhibited (**Fig. 4B**). Transformation of *comP^S34R^* was ~1 log lower than the parent when *pilTU* genes were intact, suggesting an impact of this major pilin mutation on motor-dependent dynamic activity (**Fig. 4A**). However, strikingly, transformation was readily detectable in a Δ*pilTU comP^S34R^* mutant, ~100-fold above the limit of detection in the assay, which is represented by the *ΔpilTU comP^parent^* strain (**Fig. 4A**). Importantly, this effect was not dependent on the *comP^T123C^* mutation required for pilus labeling (**Fig. S5**). These data are consistent with the model in which a bulky residue in α1C is <u>sufficient</u> to promote spontaneous disassembly during motor-independent retraction.

**Fig. 4.**
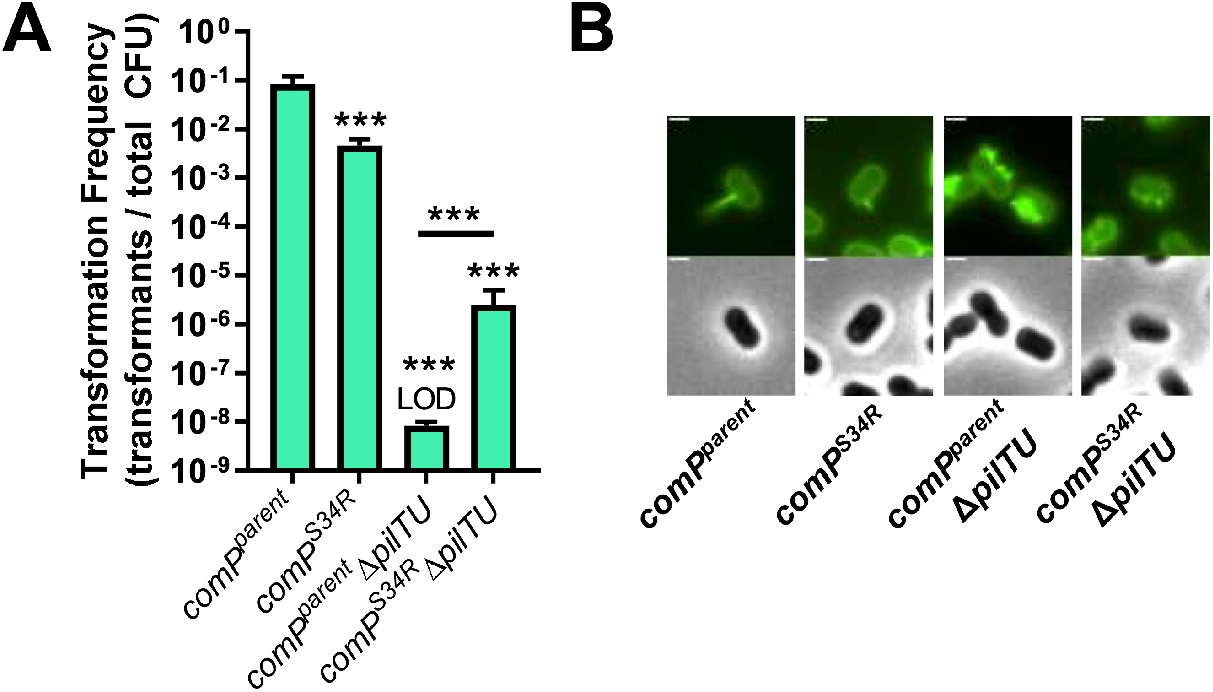
Residue bulkiness in α1C is sufficient to promote motor-independent retraction of the *A. baylyi* competence pilus. **(A)** Natural transformation assays of the indicated strains. Reactions were incubated with 50 ng of tDNA. All strains, *n* = 3. Data are shown as the mean ± SD. Asterisk(s) directly above bars denote comparisons to the parent strain. Comparisons were made by one-way ANOVA with Tukey’s post test. LOD, limit of detection; *** = *P* < 0.001. **(B)** Representative images of surface piliation for the indicated strains. Scale bar, 1 μm.

### Residue bulkiness in α1C is necessary for motor-independent retraction of the V. cholerae toxin co-regulated pilus

Next, we tested whether bulkiness in α1C was necessary for the retraction of the naturally motor-independent T4b toxin-coregulated pilus (TCP) of *V. cholerae*. TCP has no known retraction motors yet is retractile, and retraction was previously proposed to occur via spontaneous depolymerization (5). TCP dynamic pilus activity can be functionally assessed using a transducing phage (CTXΦ-Km) that confers kanamycin resistance (40–42). During transduction, CTXΦ-Km binds to the minor pilin TcpB at the pilus tip (42). TCP retraction is thought to be required for CTXΦ infection, as is the case for many phages that use T4P as receptors (5, 29, 42–45). We formally tested whether CTXΦ-Km transduction requires TCP retraction by using bulky conjugates (biotin-maleimide + neutravidin) to coat cysteine knock-in pili (*tcpA^T22C^*), which sterically obstructs pilus retraction as previously described (1, 2). Indeed, sterically obstructing pilus retraction inhibits CTXΦ-Km transduction (**Fig. S6A**). Previous work showed TCP retraction is triggered by incorporation of the minor pilin TcpB into the growing pilus, which presumably stalls pilus extension and initiates TCP retraction (5). All T4P require a glutamate at position 5 of the major pilin for pilus assembly to proceed. This glutamate forms a salt bridge with the N-terminal amine of a neighboring pilin that facilitates subunit docking and neutralizes these charges in the hydrophobic core of the pilus (35). The minor pilin TcpB also requires Glu5 to incorporate into the growing TCP to stall pilus assembly; Thus, a *tcpB^E5V^* mutant cannot initiate pilus retraction (5) and was included as an additional control. As expected, we observed that the *tcpB^E5V^* mutant exhibits dramatically reduced transduction relative to the parental strain (**Fig. 5A**), as has been observed previously (5, 46).

**Fig. 5.**
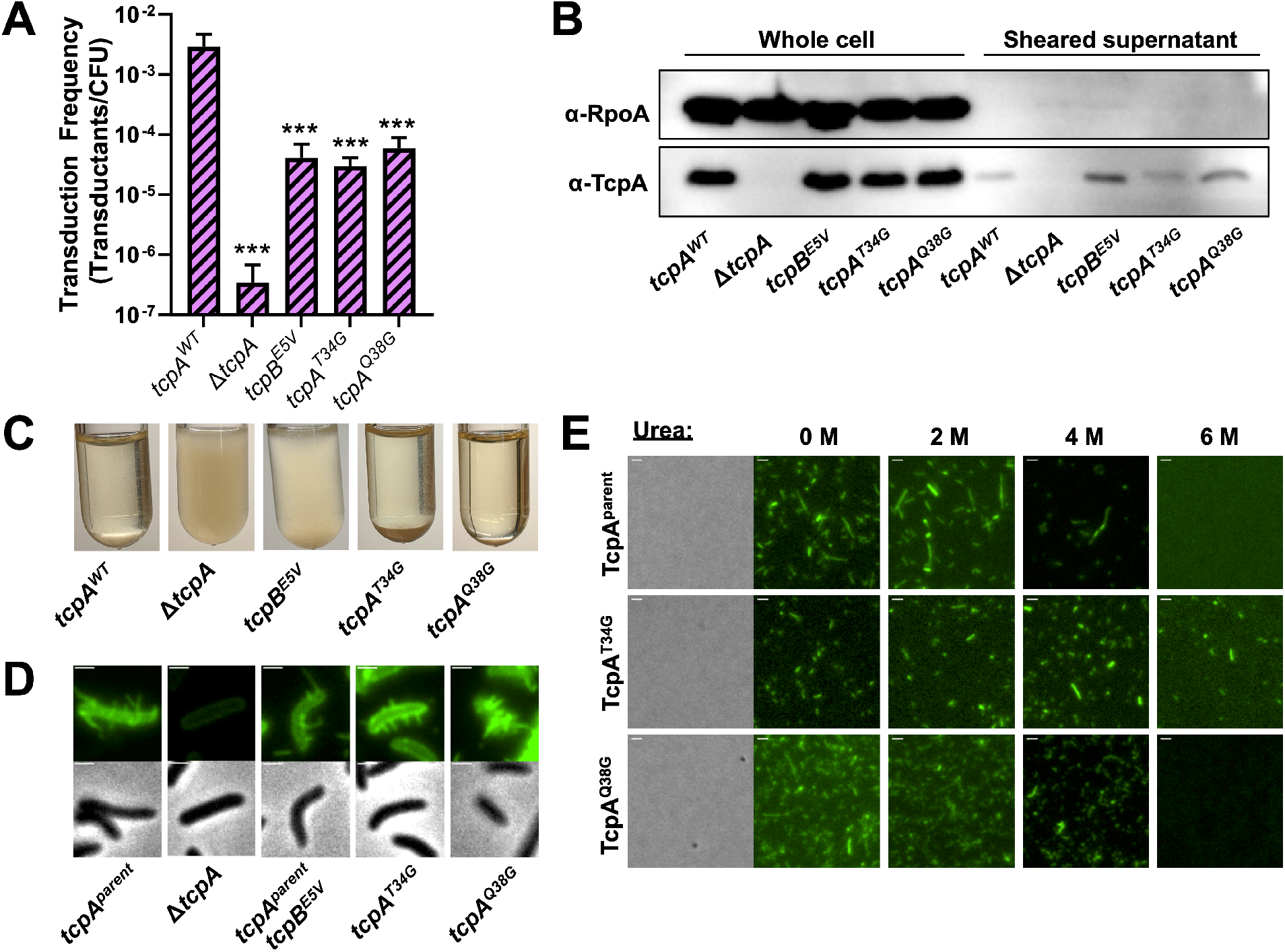
Residue bulkiness in α1C helix is necessary for motor-independent retraction in the *V. cholerae* toxin co-regulated pilus. **(A)** Transduction assays of the indicated strains. All strains, *n* = 3. Data are shown as the mean ± SD. Asterisk(s) directly above bars denote comparisons to the parent strain. Comparisons were made by one-way ANOVA with Tukey’s post test. *** = *P* < 0.001. **(B)** Western blot of the indicated strains to detect the major pilin TcpA and RpoA (as a loading and cytoplasmic protein control) in both whole cell lysates and sheared supernatant samples as indicated. Data are representative of two independent experiments. **(C)** Representative images of aggregation phenotypes and **(D)** surface piliation for the indicated strains. Scale bars in **D**, 1 μm. **(E)** Stability assay comparing chemical stability of purified AF488-mal labeled pili from the indicated strains. A representative phase-contrast image is shown to highlight that pilus preps are cell free. Fluorescent images show the presence or absence of intact pili under the indicated conditions. All images are representative of three independent biological replicates. Scale bar, 1 μm.

To test if residue bulkiness in the pilus core is necessary for TCP retraction, we targeted residues in TcpA at Thr34 (corresponding to the position of the PilA^G34R^ suppressor) and at Gln38 (a large residue one helical turn away in α1C) (**Fig. S4** and **Fig. S7)**. These bulky residues were changed to the smallest amino acid, glycine (*tcpA^T34G^*, *tcpA^Q38G^*). We hypothesized that if these bulky residues are required to destabilize the pilus and facilitate retraction, substituting them with smaller residues would reduce TCP retraction frequency and, therefore, reduce phage transduction. Indeed, both the *tcpA^T34G^* and *tcpA^Q38G^* mutants showed a significant reduction in CTXΦ transduction similar to the *tcpBE5V* strain (**Fig. 5A**). A trivial explanation for this phenotype, however, could be that these mutants prevented TCP biogenesis. To test this, we took three approaches to assess the presence of surface exposed TCP. First, TCP were sheared from the bacterial surface. Western blot analysis for TcpA in sheared surface fractions indicated that both *tcpA^T34G^* and *tcpA^Q38G^* produced similar amounts of surface associated pili as the parent and *tcpBE5V* mutant (**Fig. 5B**). Second, surface associated TCP interact, resulting in aggregation of cells in liquid culture. Aggregation of both *tcpA^T34G^* and *tcpA^Q38G^* were indistinguishable from the parent (**Fig. 5C**). Third, TCP can be labeled via the same cysteine knock-in and maleimide dye approach described above (*tcpA^T22C^*) (2, 28). We introduced these mutations into the *tcpA^T22C^* background to directly observe surface piliation by epifluorescence microscopy. Importantly, both *tcpA^T34G^* and *tcpA^Q38G^* still exhibit reduced transduction in this background (**Fig. S6B**). And consistent with our previous results (**Fig. 5B-C**), *tcpA^T34G^* and *tcpA^Q38G^* produce long external TCP filaments similar in appearance to the parent strain (**Fig. 5D**). These results indicate that the reduction in motor-independent retraction observed for the *tcpA^T34G^* and *tcpA^Q38G^* mutants is not due to reduced pilus biogenesis and are thus consistent with our hypothesis that smaller residues in α1C allow for more stable subunit packing, reducing the frequency of spontaneous retraction.

To test this hypothesis, we purified labeled TCP filaments from the parent, *tcpA^T34G^* and *tcpA^Q38G^* strains and tested their thermal and chemical stability as well as protease susceptibility as described above. Pilus filaments from all three strains have similar thermal stability and protease susceptibility profiles (**Fig. S6C**), however TcpAT34G filaments are more resistant to denaturation by 6 M urea compared to the parent and TcpAQ38G filaments (**Fig. 5E**). This is consistent with our hypothesis that small amino acids in α1C of the major pilin allow for stronger pilin-pilin interactions and increase the stability of the pilus, thereby decreasing its propensity to spontaneously retract. In addition to reducing residue bulkiness at these locations, the *tcpA^T34G^* and *tcpA^Q38G^* mutations also modify the polarity at this position (a change from polar to non-polar amino acids). For these studies with TCP, we cannot rule out that destabilization occurs not from the relief of steric stress in the TcpA^T34G^ mutant, but from loss of polar interactions between Thr34 and the α1N of adjacent pilin subunits. Together, these data further support the idea that bulky residues in α1C oriented into the pilus core are <u>necessary</u> to promote motor-independent retraction.

### The presence of bulky residues in α1C is a conserved property of retraction motor-independent pilus systems

Because we found that bulky residues in α1C were necessary for motor-independent retraction in three distinct T4P, we next sought to address whether this was a conserved feature of retraction motor-independent systems. Specifically, we hypothesized that retraction motor-independent systems would have bulkier residues in α1C that destabilize filaments, facilitating spontaneous depolymerization; whereas retraction motor-dependent systems may tolerate smaller residues in this region that fit better into the filament core, resulting in more stable pili that do not permit spontaneous depolymerization. T4P fit into the broader family of T4F, which includes several closely related systems that appear to retract without a dedicated motor, such as the Gram-positive Com pili and the type II secretion system (T2SS) (20, 47–50). T2SSs are widely conserved in Gram-negative bacteria and work through the action of a putatively dynamic pseudopilus for export of proteins across the outer membrane (51, 52). We predicted that if the T4Fs from these systems also retract by spontaneous depolymerization, as has been previously suggested (5, 36), that they might also have bulkier residues in α1C that destabilize the filament.

To investigate this, we aligned the α1 region of major pilins from motor-dependent T4P, motor-independent T4P, Com pili, and T2SSs (**Fig. S7C**). The residues of α1C that are oriented into the pilus core were determined based on structural models of the *V. cholerae* PilA, *A. baylyi* ComP, and *V. cholerae* TcpA major pilins (**Fig. S7B**). To highlight differences between bulky residues in this region, we grouped amino acids into four categories based on the size of their R-group (**Fig. S7A**), mapped these designations onto the alignments (**Fig. S7B**) and plotted them (**Fig. 6A**) to compare T4F major pilins. We found that major pilins from retraction motor-independent T4P, Com pili, and T2SSs generally have bulkier residues that protrude into the pilus core compared to major pilins from diverse retraction motor-dependent pilus systems (**Fig. 6A**). Interestingly, bulkiness at the 34 position in α1C, which is the predominant position we have studied by mutational analysis, did not exhibit a clear trend when comparing motor-dependent and motor-independent T4Fs (**Fig. S7C**). Experimental introduction of bulkiness at this site, however, slightly diminished function of the *A. baylyi* competence T4aP (ComP^S34R^) (**Fig. 4A**), and reduced pilus length of the *V. cholerae* competence T4aP (PilA^G34R^) (**Fig. 2B**). Thus, while bulky residues at this site may facilitate motor-independent retraction, they may also exhibit pleiotropic effects on other pilus properties. In TCP, we show that a bulky residue in α1C one helical turn away from position 34 also contributed to motor-independent retraction (TcpA^Q38^) (**Fig. 5**). Thus, it may be that the cumulative bulkiness of core-oriented residues in the rigid α1C are critical for destabilizing the subunit interactions, promoting motor-independent retraction. When we compare the number of small residues in α1C that are core-oriented, we find that retraction motor-independent T4P, Com pili, and T2SSs have significantly fewer small residues within this region compared to retraction motor-dependent T4P (**Fig. 6B**), consistent with bulkiness in core-oriented α1C residues being a conserved feature among motor-independent T4Fs.

**Fig. 6.**
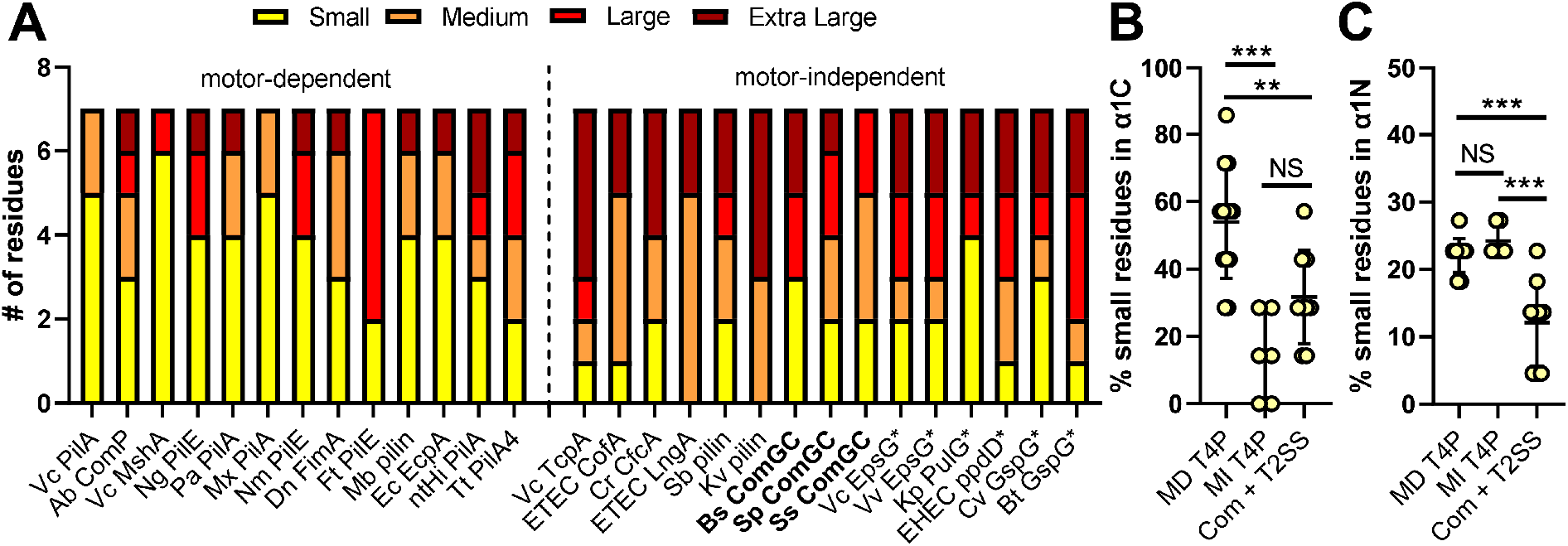
Major pilins from motor-independent T4Fs generally contain more bulky core-oriented residues in α1C. **(A)** Stacked bar chart showing the number of amino acids that protrude into the pilus core in the α1C domain that are small, medium, large or extra large. See **Fig. S7A-C** for details on how these data were generated. **(B)** From **A,** the percent of small residues in α1C that are core-oriented. **(C)** The percent of small residues in the α1N region that likely interact with α1C of neighboring subunits (residues 2-22). **B** and **C** are plotted to compare the major pilins of retraction motor-dependent (MD) T4P, retraction motor-independent (MI) T4P, and Com pili + T2SSs. Motor-dependent pilus systems are defined as requiring a retraction motor for normal pilus activity and/or having a predicted retraction ATPase gene in the pilus operon. Motor-independent pilus systems are defined as those that lack canonical retraction ATPase genes. Data are shown as the mean ± SD and statistical comparisons are made one-way ANOVA with Tukey’s post test. NS, not significant; ** = *P* < 0.01, *** = *P* < 0.001.

Our hypothesis is that bulky residues in α1C destabilize pili and promote spontaneous depolymerization by disrupting interactions between this α1C region and α1Ns of adjacent pilins. However, large residues in α1C could be compensated with small residues in α1N to maintain pilus stability. If true, we expected an inverse trend in residue bulkiness for α1N when comparing motor-dependent and motor-independent T4Fs. Specifically, we assessed residue bulkiness in the α1N region that likely interacts with the α1C of neighboring subunits (residues 2-22) (**Fig. S7C**). This analysis revealed no difference in the general trend for α1N bulkiness between the motor-dependent T4P and the motor-independent T4Fs (**Fig. 6C**). Together, these data suggest that retraction motor-independent T4Fs have bulkier residues in α1C as a conserved feature, which may act to destabilize the pilus filament and allow for retraction by spontaneous depolymerization.

## DISCUSSION

This study sheds light on the mechanisms underlying pilus retraction by providing evidence to support a long-standing model in which motor-independent retraction occurs via the spontaneous depolymerization of the pilus filament (5, 36, 37). In particular, we reveal that the stability of the pilus filament is a critical determinant of motor-independent retraction in diverse T4P, including systems that harbor retraction motors (the competence pili of *V. cholerae* and *A. baylyi*) as well those that lack a retraction motor (the toxin co-regulated pili of *V. cholerae*). Notably, we demonstrate that bulky residues oriented toward the pilus core in α1C of the major pilin, which may destabilize pilin-pilin interactions, are both necessary and sufficient to facilitate motor-independent retraction in these three distinct T4P. Thus, our results uncover a broadly conserved mechanism for T4P retraction that is inherent to the pilus filament.

Spontaneous depolymerization is proposed to occur when pilus assembly stalls, either by incorporation of the minor pilin TcpB in the case of *V. cholerae* TCP (5), or perhaps due to dissociation of the extension ATPase (36). Upon stalling, the terminal pilin subunit at the base of the growing pilus is in a state of dynamic instability where it can diffuse back into the inner membrane (**Fig. 7**) (5, 36). Loss of this terminal pilin will cause the pilus to collapse into the inner membrane, positioning the next subunit into this unstable state, and resulting in the iterative spontaneous depolymerization of the pilus filament. This model describes a Brownian ratchet whereby the extended pilus possesses potential energy from the ATPase activity of the assembly motor, but defaults to a lower energy state with release of each subunit back into the membrane after assembly stalls (37). We propose that inter-subunit contacts between the terminal pilin and the closest pilin subunits at the base of the filament represent critical contacts that determine whether or not a pilus will depolymerize spontaneously. Strong interactions between subunits reduce the likelihood of spontaneous depolymerization, stabilizing a non-retracting extended pilus filament (**Fig. 7**; yellow pilin). On the other hand, weaker pilin-pilin contacts enhance the likelihood that the terminal pilin will diffuse away from the base of a stalled pilus to trigger spontaneous depolymerization (**Fig. 7**; red pilin). This may explain the increase in retraction frequency observed in our Δ*pilTU pilA* suppressor mutants. Furthermore, residues 34, 78 and 80 in PilA all lie near the bottom of the pilin globular domain that presumably sits on the inner membrane. Disrupting the contacts that these residues make with α1N of neighboring subunits may help to release this subunit from the base of the pilus if assembly is stalled (**Fig. 7**). Thus, pilin-pilin interactions between the terminal pilin and the closest pilin subunits at the base of the filament may represent the most critical determinant for retraction by spontaneous depolymerization. We have characterized one key interface—core-oriented residues in the α1C—that contribute to this process, however, other interaction interfaces within the filament may also play a critical role. In addition to the pilin interactions in the pilus filament, the inner membrane platform protein may also influence the tendency of a subunit to diffuse back into the membrane; subunits in motor-dependent retraction T4P may be stabilized by the platform protein in a motor-dependent manner to prevent spontaneous diffusion back into the membrane (36).

**Fig. 7.**
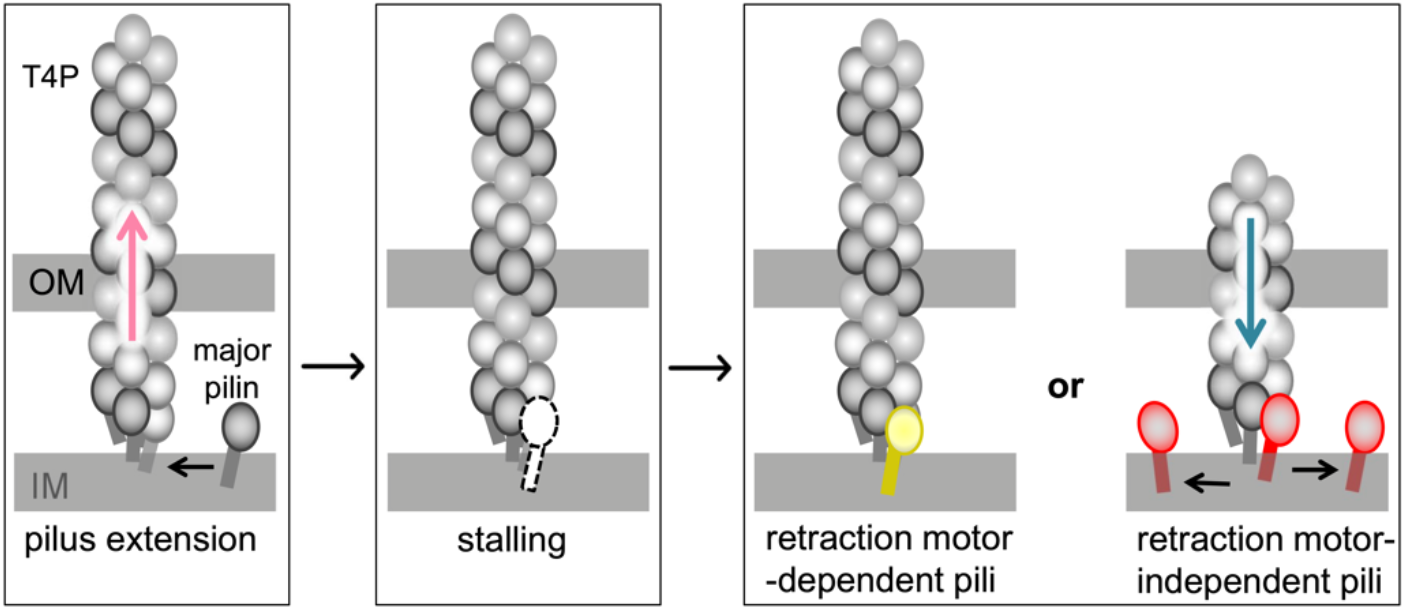
Model for motor-independent retraction of T4P. Pilus polymerization involves docking of the major pilin at the pilus base followed by ATP hydrolysis by the assembly motor and outward extrusion of the growing filament. When pilus assembly stalls, the terminal pilin subunit enters a state of dynamic instability (white pilin with dashes). In retraction motor-dependent pili, pilin-pilin interactions are strong (yellow pilin), and the terminal pilin is more stably associated with the filament, which may prevent its diffusion back into the membrane. In retraction motor-independent pili, pilin-pilin interactions are sufficiently weak (red pilin), and the terminal pilin is less stably associated with the base of the filament, which may favor its diffusion into the membrane. Loss of this terminal pilin will cause the pilus to collapse into the inner membrane, positioning the next subunit into this unstable state. The iterative diffusion of the terminal pilin from the base of the filament results in motor-independent retraction via spontaneous depolymerization. One factor involved in destabilizing pilin-pilin interactions may be bulkiness of residues in α1C that are oriented into the pilus core. If residues in this region are small (yellow pilin), the resulting pilin-pilin interactions are strong and the ability of the filament to retract in the absence of motors is reduced. Conversely, if residues in α1C are large (red pilin), the resulting pilin-pilin interactions may be weaker and the filaments are more likely to undergo motor-independent retraction.

Differences in the dynamics of the terminal pilin at the base of the pilus are reflected in the overall stability of the pilus filament. Using mutants of both *V. cholerae* motor-dependent competence pili and motor-independent TCP, we demonstrate that filament stability correlates with motor-independent retraction where mutants with reduced filament stability are more likely to retract, and mutants with increased filament stability are less likely to retract. Retraction motor-dependent gonococcal pili are more stable than retraction motor-independent TCP, a feature that was attributed, in part, to the presence of several stacking aromatic residues in gonococcal pili that are absent in TCP (38). Here we find that the retraction motor-dependent competence T4aP in *V. cholerae* are also more stable than TCP, but that stability can be altered with a single amino acid change. Generally, retraction motor-dependent T4aP retract with significantly higher forces compared to retraction motor-independent pilus systems (1, 5, 23, 53). The high retraction force of many T4aP is likely essential for their functions such as twitching motility and DNA uptake. Thus, it has been proposed that T4aP have evolved more stable filaments to withstand the high retraction forces they experience when attached to a load such as an epithelial cell or DNA (38). Based on our results, however, it is tempting to speculate that filament instability is an evolutionarily maintained property of naturally retraction motor-independent pilus systems because the poor pilin-pilin interactions that cause this instability are essential for these pili to retract by spontaneous depolymerization. Our data also show that reducing pilus filament stability does not impact motor-dependent pilus retraction (**Fig. S1**). This suggests that T4aP may have gained the ability to evolve more stable filaments because the acquisition of retraction motors allowed these systems to overcome the tradeoff that would have prevented these pili from retracting by spontaneous depolymerization (36).

As mentioned above, the T4cP of *C. crescentus* utilize a bifunctional ATPase motor, CpaF, to power both pilus extension and retraction. Our data indicate that for *V. cholerae* competence T4aP, the PilB ATPase is not a bifunctional motor (**Fig. 1C**). T4cP, however, are quite divergent from the T4aP/T4bP, and represent a distinct evolutionary branch (2, 20, 38, 49). T4cP have also evolved two distinct platform proteins, a property that is distinct from the T4aP and T4bP, which may be required for the activity of a bifunctional motor (20, 29). Therefore, T4cP and T4aP/T4bP may have evolved distinct mechanisms for retraction. T4aP represent a relatively recent branch of the T4bP and we imagine that the T4aP that exhibit pilus retraction in the absence of dedicated retraction motors may have maintained the residual ability to spontaneously depolymerize (2, 49). Both Com pili and T2SSs are more closely related to the T4aP/T4bP than the T4cP (2, 49). Based on our findings, we hypothesize that motor-independent retraction via spontaneous depolymerization may be a broadly conserved property of T4Fs. However, since the vast majority of these systems are untested, it remains possible that retraction of some T4Fs may be supported by a bifunctional motor or an as yet undiscovered mechanism.

## ACKNOWLEDGEMENTS

We would like to thank Nicolas Biais and Stefan Kreida for helpful discussions; Kim D. Seed and Zoe Netter for protocols and strains for transduction assays; Courtney K. Ellison for providing protocols for pilus labeling and purification; and Triana N. Dalia for help with strain construction. This work was supported by Grant R35GM128674 from the National Institutes of Health (to ABD), and Grant RGPIN-2017-05757 from the Natural Sciences and Engineering Research Council (to LC).

## MATERIALS AND METHODS

### Bacterial strains and culture conditions

*Vibrio cholerae* and *Acinetobacter baylyi* strains were routinely grown in LB Miller broth and on LB Miller agar supplemented with erythromycin (10 μg/mL), kanamycin (50 μg/mL) and spectinomycin (50 μg/mL or 200 μg/mL) when appropriate.

### Construction of mutant strains

*V. cholerae* strains used throughout this study are derivatives of the El Tor isolate E7946. All *A. baylyi* strains used are derivatives of strain ADP1 (54). Throughout the study, strains with or without cysteine knock-ins in the major pilin of the indicated T4P were used depending on the experimental approach. When the cysteine knockin background was used, the parent strain is denoted as *pilA^parent^*, *comP^parent^*, or *tcpA^parent^* (open bar graphs). If the non-Cys background was used (i.e. the pilin sequence was wild type), then the parent strain is denoted as *pilA^WT^*, *comP^WT^*, or *tcpA^WT^* (hashed bar graphs). All mutant constructs were generated by splicing-by-overlap extension (SOE) PCR and then introduced into strains via MuGENT and natural transformation exactly as previously described (31, 55–57). Amino acid positions are numbered according to the mature pilin sequence. A list of all the mature pilin residue positions mentioned in the text and their corresponding positions relative to their start codon are listed in **Table S1**. For a detailed list of all mutant strains used throughout this study see **Table S2**. For a detailed list of all primers used to construct mutant strains see **Table S3**and for a list of the genes mutated in this study see **Table S4**.

### Natural transformation assays

In order to induce maximal competence in *V. cholerae* strains, the master competence regulator TfoX is overexpressed using an IPTG-inducible P_*tac*_ promoter and the cells are genetically locked in a regulatory state that mimics high cell density via deletion of *luxO,* as previously described (56, 58–62). Chitin-independent transformation assays were performed exactly as previously described (31). Briefly, strains were grown overnight with aeration at 30 °C with 100 μM IPTG then ~10^8^ colony forming units (CFU) were subcultured into 3 mL of LB + 100 μM IPTG + 20 mM MgCl_2_ + 10 mM CaCl_2_ and grown to late log phase. Next, ~10^7^ CFU of this culture was diluted into Instant Ocean medium (IO) (7 g/L; Aquarium Systems) supplemented with 100 μM IPTG and 200 ng of transforming DNA (tDNA) was added to each reaction and incubated statically at 30 °C overnight. The tDNA replaces VC1807, a frame-shifted transposase, with an erythromycin resistance antibiotic marker (aka ΔVC1807::Erm^R^) as previously described (55). Negative control reactions where no tDNA was added were performed for each strain. After incubation with tDNA, reactions were outgrown by adding 1mL of LB to each reaction and shaking (250 rpm) at 37 °C for ~2 hours. Reactions were then plated for quantitative culture onto agar plates selecting for transformants (LB + 10 μg/mL erythromycin) or onto plain LB for total viable counts. The transformation frequency is defined as the number of transformants divided by the total viable counts. For reactions where no transformants were obtained, a limit of detection was calculated and plotted.

To physically obstruct pilus retraction, ~10^8^ CFU of the indicated strains grown to late log phase under competence-inducing conditions were washed in IO and incubated with 50 μg/mL biotin-maleimide (biotin-mal, Thermo Fisher) for 45 min at room temperature. Then, cells were pelleted and washed twice in IO. Next, ~10^7^ CFU were diluted into fresh IO. Where indicated, neutravidin (Thermo Fisher) was added to reactions at a final concentration of 1.32 mg/mL and incubated at room temperature for 30 min. Then, 200 ng ΔVC1807::Erm^R^ transforming DNA was added and reactions were incubated at 30 °C for 10 minutes before the addition of 10 units of DNAse I (NEB) to prevent additional DNA uptake. This was performed because the production of new pilins during the normal 5h tDNA incubation would circumvent our method to block pilus retraction. Following DNaseI addition, all reactions were incubated at 30 °C for a total of 5 h to allow for DNA integration. Reactions were then outgrown and plated exactly as described above to attain the transformation frequency.

For *A. baylyi*, transformations were performed exactly as previously described (63). Competence in *A. baylyi* is constitutively active. Briefly, strains were grown overnight in LB medium. Then, ∼10^8^ cells were diluted into fresh LB medium, and tDNA was added (∼50 ng). Reactions were incubated at 30°C with agitation for 5 hours and then plated for quantitative culture as described above to determine the transformation frequency. The tDNA targeted ACIAD1551 for deletion and replacement with an Ab^R^ as previously described (aka ΔACIAD1551::Ab^R^) (63). Negative control reactions where no tDNA was added were also performed for each strain.

### Pilin labelling, imaging and quantification

In order to label *V. cholerae* competence pili for observation with epifluorescence microscopy, strains were grown to late log exactly as described above for natural transformation assays. Then, ~10^8^ CFU were spun down at 18,000 x g for 1 min and resuspended in IO. Cells were then incubated with 25 μg/mL AF488-mal for 15 minutes statically at room temperature in the dark. Cells were washed twice, fully disrupting the pellet using 100 μL of IO for each wash, and finally resuspended in IO to a final concentration of ~10^7^ CFU/mL.

In order to label *V. cholerae* toxin co-regulated pili (TCP), virulence was induced by overexpressing the master competence regulator ToxT using an IPTG-inducible P_*tac*_ promoter. For observation with epifluorescence microscopy, strains were grown to late log in LB + 10 μM IPTG. Then, ~10^8^ CFU were directly incubated with 25 μg/mL AF488-mal for 5 minutes statically at room temperature in the dark. Cells were then spun down at 5,000 x g for 3 minutes and the resulting pellet was gently washed (without disrupting the pellet) three times with 100μL IO. Cells were then resuspended to a final concentration of ~10^7^ CFU/mL.

In order to label *A. baylyi* competence pili for observation, strains were grown to mid log in LB. ~108 CFU were spun down at 18,000 x g for 1 min and resuspended in LB. then cells were incubated with 25 μg/mL AF488-mal for 15 minutes statically at room temperature in the dark. Labeled cells were pelleted, gently washed once with 100 μl of LB without disrupting the pellet, and resuspended in 50 μl 1X PBS [10mM Na_2_HPO_4_, 1.8mM KH_2_PO_4_, 2.7mM KCl, 137mM NaCl].

For imaging, 2μL of labeled cells were placed under an 0.2% gelzan pad (made in IO for *V. cholerae* or 1X PBS for *A. baylyi*) on a coverslip and imaged. All imaging was done on an inverted Nikon Ti-2 microscope with a Plan Apo ×60 objective, a green fluorescent protein filter cube, a Hamamatsu ORCAFlash 4.0 camera and Nikon NIS Elements imaging software. Representative images of the piliation state of strains were gathered and unless otherwise noted, the lookup tables for each phase or fluorescent image were adjusted to the same range in each figure.

In order to measure extension and retraction events, labelled cells were imaged by time-lapse microscopy. A phase-contrast image (to image cell bodies) and fluorescent image (to image labeled pili) were taken every second for 1 minute or every 10 seconds for 10 minutes. To enhance the dynamic activity of Δ*pilTU* strains, a tightly inducible construct of *pilB* (P_*BAD*_-*pilB*) was used in a Δ*pilB* background. These strains were grown to late log without inducer and then 0.2% arabinose was added to cells 30 minutes before imaging. Cells were then imaged under a 0.2% IO gelzan pad infused with 0.2% arabinose. Rates were calculated manually using measurement tools in the NIS Elements analysis software. Only pili longer than 0.3 μm and that completed extending or retracting within the time-lapse window were analyzed.

In order to calculate the retraction frequency, time-lapse videos of cell bodies and labeled pili were captured using phase-contrast and epifluorescence microscopy, respectively. Time-lapses from 3 independent biological replicates were sectioned into areas containing ~100-500 cells. The number of cells that exhibited a pilus retraction event and the total number of cells in the field were manually counted.

To assess aggregation and pelleting of cultures, strains were grown to late log under either competence or virulence inducing conditions as described above. Cultures were then allowed to sit statically at room temperature for 30-60 mins. Images of culture tubes were then taken against a white background.

### Western Blotting

Strains were grown to late log in LB with 10 μM IPTG to induce production of TCP as stated above. For whole cell samples, 1x Fastbreak Lysis Buffer (Promega), 38 μg/mL lysozyme (Gold Bio), and 25 units Benzonase (Sigma) were added to 100 μL of culture and incubated at RT for 10 minutes. For sheared supernatant samples, 100 μL of culture was vigorously vortexed for 2 minutes. Samples were then centrifuged at 5,000 x g for 10 minutes to pellet cell bodies and the supernatant containing sheared pili was carefully removed. Resulting whole cell and sheared supernatant samples were then mixed 1:1 with 2X SDS sample buffer [220 mM Tris pH 6.8, 25% glycerol, 1.8% SDS, 0.02% Bromophenol Blue, 5% *β*-mercaptoethanol] and then boiled for 10 minutes @100 °C. Then 10 μL of each sample was separated on a 15% SDS PAGE gel. Proteins were electrophoretically transferred to a PVDF membrane and incubated with both of the following primary antibodies: α-TcpA polyclonal rabbit and α-RpoA monoclonal mouse (Biolegend). Then, blots were washed and incubated with a α-rabbit and α-mouse secondary antibodies conjugated to horseradish peroxidase (HRP) and developed using Pierce ECL Western blotting substrate. Blots were then imaged using a ProteinSimple FluorChem E instrument.

### Transduction Assay

Cholera toxin phage harboring kanamycin resistance (CTXΦ-Km) was isolated from *V. cholerae* donor strains containing Δ*ctxAB*::KmR and *lexA*, the SOS response repressor gene, under the control of tightly inducible P_*BAD*_-riboswitch promoter (64). CTXΦ excision from the genome occurs when the SOS response is activated so regulating the expression of *lexA* allows for the controlled induction of the SOS response and CTXΦ production. CTXΦ-Km phage was prepared by growing the donor strain in LB + 0.2% arabinose +1.5 mM theophylline overnight with aeration at 30 °C. Cells were washed twice with LB by centrifugation at 18,000 x g for 1 minute and 10 μL of washed culture was used to inoculate 3 mL of plain LB (i.e. without inducers to allow for depletion of LexA) and grown at 30°C with aeration for 5 hours. Donor supernatant was cleared twice by centrifugation at 5,000 x g for 3 minutes to remove cells. The supernatant containing CTXΦ-Km was then flash frozen and stored at −80°C and thawed on ice when needed.

The production of TCP was induced in *V. cholerae* recipient strains using a P_*tac*_-*toxT* construct as described above. These strains also harbor a deletion of *ctxAB* to attenuate these strains. Recipients are SpecR, while the CTXΦ-Km donor strain is sensitive to spectinomycin. So, spectinomycin was used to counter-select any contaminating donor strain that might still be present in the donor phage supernatant. For preparation of recipient strains, cultures were diluted to OD_600_ = 0.05 in LB and grown shaking at 37 °C to OD_600_ = 0.3 (41). Then, cultures were incubated statically at 37 °C for 2 hours after addition of 10 mM MgCl_2_ and 10 μM IPTG. Following this, donor phage supernatant and recipient culture were mixed together in a 1:3 ratio and incubated for 1 hour statically at 37 °C. Reactions were then plated for quantitative culture onto agar plates selecting for transductants (LB + 200 μg/mL spectinomycin + 50 μg/mL kanamycin) and onto LB + 200 μg/mL spectinomycin for total viable counts. The transduction frequency is defined as the number of transductants divided by the total viable counts. Negative control reactions where no donor supernatant was added to the recipient were also performed for each strain.

To physically obstruct TCP retraction, ~10^8^ CFU of the recipient strains prepared as described above were pelleted at 5,000 x g for 3 minutes, resuspended in IO and incubated with 50 μg/mL biotin-mal for 45 min at room temperature. Then, cells were pelleted and gently washed in IO twice without disrupting the pellet. Next, ~10^7^ CFU were diluted into fresh IO and where indicated, neutravidin was then added to reactions at a final concentration of 1.32 mg/mL and incubated at room temperature for 30 min. Then, donor CTXΦ-Km supernatant was mixed with recipient and transduction assay proceeded exactly as described above.

### Pilus filament stability assay

For purification of competence pili and TCP, the relevant *V. cholerae* strains were grown and labelled with AF488-mal dye exactly as described above, with the exception that cells were resuspended after washing to ~10^10^ CFU/mL in 25 mM Tris pH 8.8. Cells were then loaded onto a SpinX 0.2 μm filter column (Sigma) and centrifuged at 16,000 x g for 1 minute and the flow through containing pili was collected. For competence pili, the flow through was centrifuged at 10,000 x g for 10 min at 4oC to remove cellular debris.

To assess thermal stability, TCP and competence pili were incubated at RT, 40 °C, 50 °C and 60 °C for 1 hour. To assess chemical stability, pili were incubated with 0 M, 2 M, 4 M or 6 M urea for 1 hour at RT. To assess protease susceptibility, pili were mixed with proteinase K at a final concentration of 5 mg/mL and incubated at 37°C for 1 hour. Following these treatments, pili were imaged under a 0.2% IO gelzan pad as described above. Due to variable levels of background signal, the look up tables (LUTs) for TCP filament images were adjusted to the same range within a given strain for each condition.

### Protein structure image rendering

Structural renderings of major pilins were generated using PyMOL v. 2.3.4. The GenBank accession numbers for the primary sequence of major pilins and the Protein Data Bank (PDB) accession numbers of either predicted or solved structural renderings of major pilins can be found in the figure legend of **Fig. 1**, **S5** and **S6**. The Phyre2 web portal was used to generate all predicted protein structures. Residues in *V. cholerae* PilA that are “core-oriented” were chosen based on their location in the pilus filament model in **Fig 1F**, and “core-oriented” residues in the other major pilins were chosen based on their structural alignment with the Vc PilA model and their position within α1C relative to the C-terminal head group.

In order to create the model of the *V. cholerae* competence pilus filament in **Fig 1F**, the Phyre-predicted Vc PilA structure was superimposed onto one chain of the *P*. *aeruginosa* strain K (PaK) pilus filament cryoEM reonstruction (PDB 5VXY) using Chimera. To account for the melted segment of α1N that is apparent in the cryoEM reconstruction, residues 1-28 of Vc PilA were replaced with the corresponding segment of PaK pilin using the coordinates for the cryoEM reconstruction. The helical symmetry parameters for the PaK pilus (10.5 Å rise, 87.3 ° rotation) were imposed on the new Vc PilA model to generate a pilus with 19 subunits.

### Statistics

Statistical differences were assessed by Student’s t-test or one-way ANOVA tests followed by either a Dunnett’s or Tukey’s multiple comparisons post-test as indicated using GraphPad Prism software v. 9.0.0. For all transformation frequency, frequency of pilus retraction, and transduction frequency experiments statistical analyses were performed on log-transformed data. All statistical comparisons can be found in **Table S5**.

## Supplemental Information

**Fig. S1.**
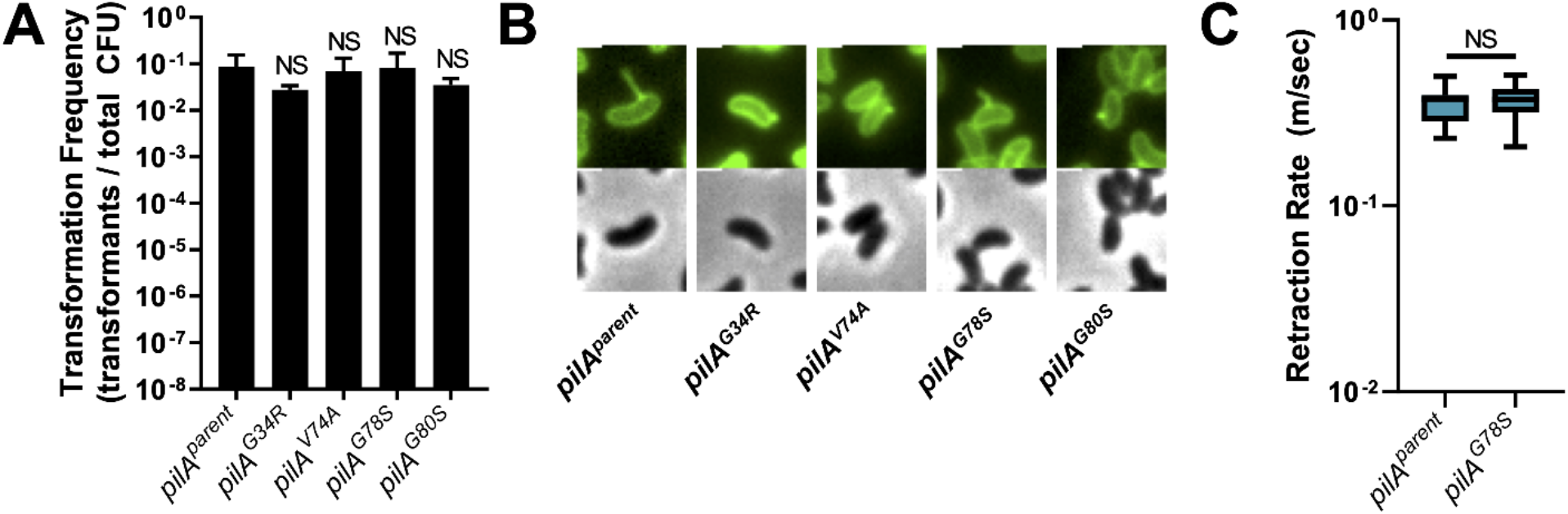
*pilA* suppressor mutations do not affect motor-dependent retraction. All strains used in this figure have *pilTU* intact to assess motor-dependent retraction. **(A)** Natural transformation assays of the indicated strains. Reactions were incubated with 200 ng of tDNA. All strains, *n* = 3. Data are shown as the mean ± SD. Notations directly above bars denote comparisons to the parent strain. **(B)** Representative images of surface piliation (top) and aggregation phenotypes (bottom) for the indicated strains. Scale bar, 1 μm. **(C)** Rates of pilus retraction were measured in the indicated strains by epifluorescence time-lapse microscopy of AF488-mal labeled cells. To enhance dynamic activity, *pilB* alleles were placed under the control of a tightly inducible P_*BAD*_ promoter in a Δ*pilB* background and the induction of *pilB* expression was delayed until just before imaging was performed. Data are from three independent biological replicates; All strains, *n* = 44. Box plots represent the median and the upper and lower quartile, while the whiskers demarcate the range. All comparisons were made by one-way ANOVA with Tukey’s post test. NS, not significant.

**Fig. S2.**
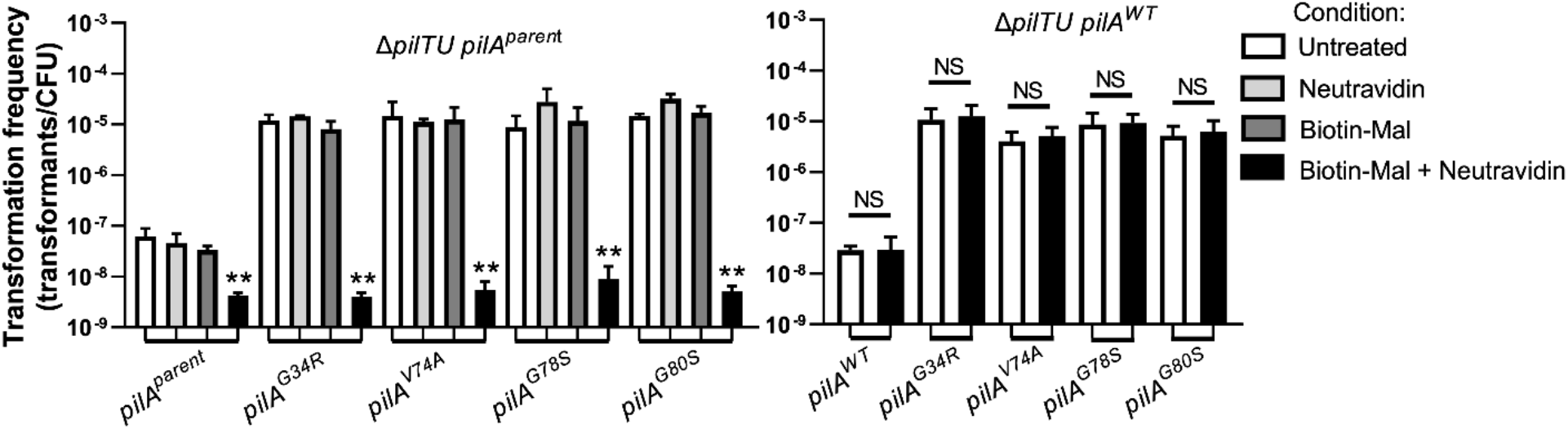
*pilA* suppressor mutants require pilus retraction to promote DNA uptake during natural transformation. Natural transformation assays of strains with *pilA^S56C^* mutation (left graph) or without the cysteine knock-in (right graph) after the indicated treatment. Reactions were incubated with 200 ng of tDNA for 10 minutes after which DNaseI was added to prevent additional DNA uptake. All strains, *n* = 3. Bar graphs show the mean ± SD. Asterisk(s) directly above bars denote comparisons to the parent strain. All comparisons were made by one-way ANOVA with Dunnett’s post test (left graph) or Tukey’s post test (right graph). NS, not significant; ** = *P* < 0.01.

**Fig. S3.**
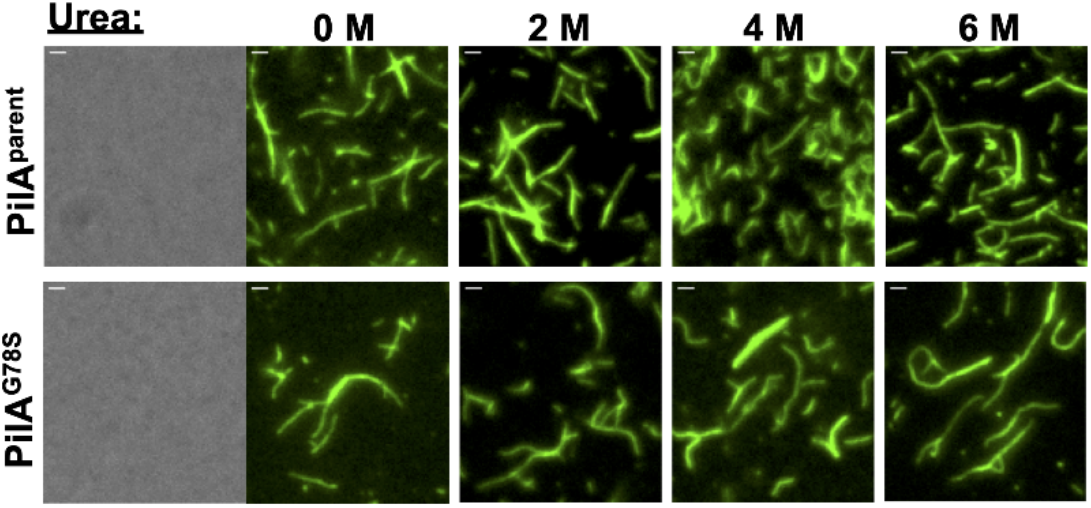
PilA^parent^ and PilA^G78S^ pilus fibers have similar chemical stability. Stability assay comparing chemical stability of purified AF488-mal labeled pilus filaments from Δ*pilTU* strains expressing the indicated *pilA* allele. A representative phase-contrast image is shown to highlight that pilus preps are cell free. Fluorescent images show the presence or absence of intact pili under the indicated conditions. All images are representative of three independent biological replicates. Scale bar, 1 μm.

**Fig. S4.**
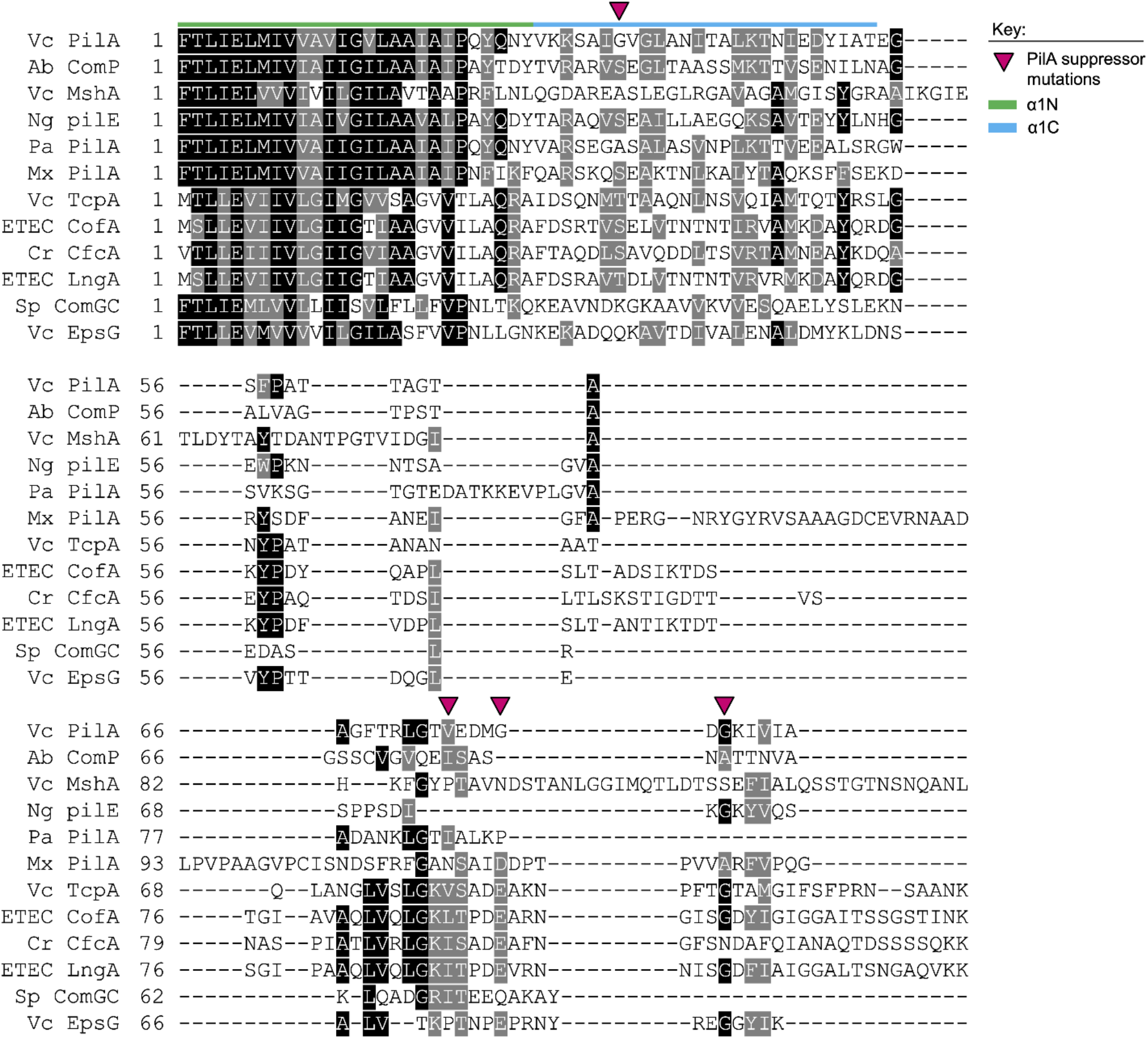
Alignment of mature pilin sequences from different pilus systems. The green (N terminal) and blue (C terminal) lines denote the α1 helix and magenta arrows indicate the position of the *pilA* suppressor mutations (*pilA^G34^*, *pilA^V74A^*, *pilA^G78S^*, *pilA^G80S^*). Residues highlighted in black are identical, while those highlighted in gray are similar. The GenBank Accession numbers for the protein sequences used are as follows: Vc, *Vibrio cholerae* PilA (NP_232053); Ab, *Acinetobacter baylyi* ComP (WP_004923779); Vc, *V. cholerae* MshA (AAF93582); Ng, *Neisseria gonorrhoeae* PilE (AAA53145); Pa, *Pseudomonas aeruginosa* PilA (WP_058135760); Mx, *Myxococcus xanthus* strain PilA (WP_011555734); Vc, *V. cholerae* TcpA (WP_001176374); ETEC, enterotoxigenic *E. coli* CofA (WP_110409206); Cr, *Citrobacter rodentium* CfcA (AAO17805); ETEC, enterotoxigenic *E. coli* LngA (AAC33154); Sp, *Streptococcus pneumoniae* ComGC (WP_054380926); Vc, *V. cholerae* EpsG (WP_000738789.1). Residue numbering in this alignment is based on the mature (processed) pilin protein.

**Fig. S5.**
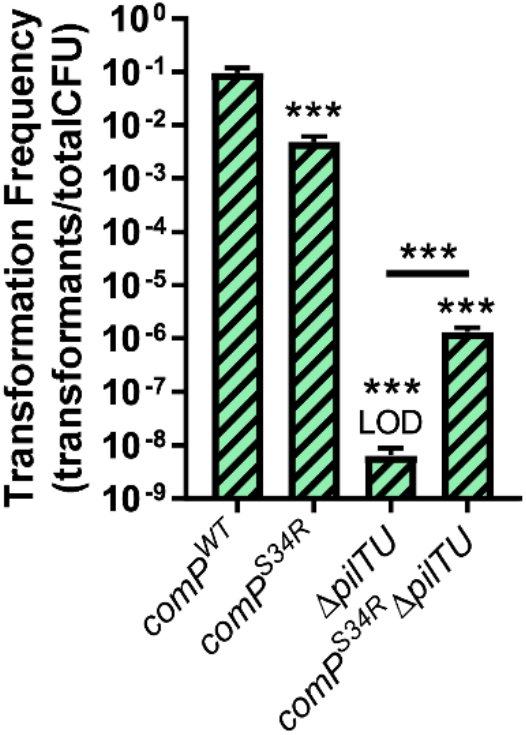
Residue bulkiness in the α1C helix is sufficient to promote motor-independent retraction of the *A. baylyi* competence pilus. **(A)** Natural transformation assays of the indicated strains. Reactions were incubated with 50 ng of tDNA. All strains, *n* = 3. Data are shown as the mean ± SD. Asterisk(s) directly above bars denote comparisons to the parent strain. Comparisons were made by one-way ANOVA with Tukey’s post test. LOD, limit of detection; *** = *P* < 0.001.

**Fig. S6.**
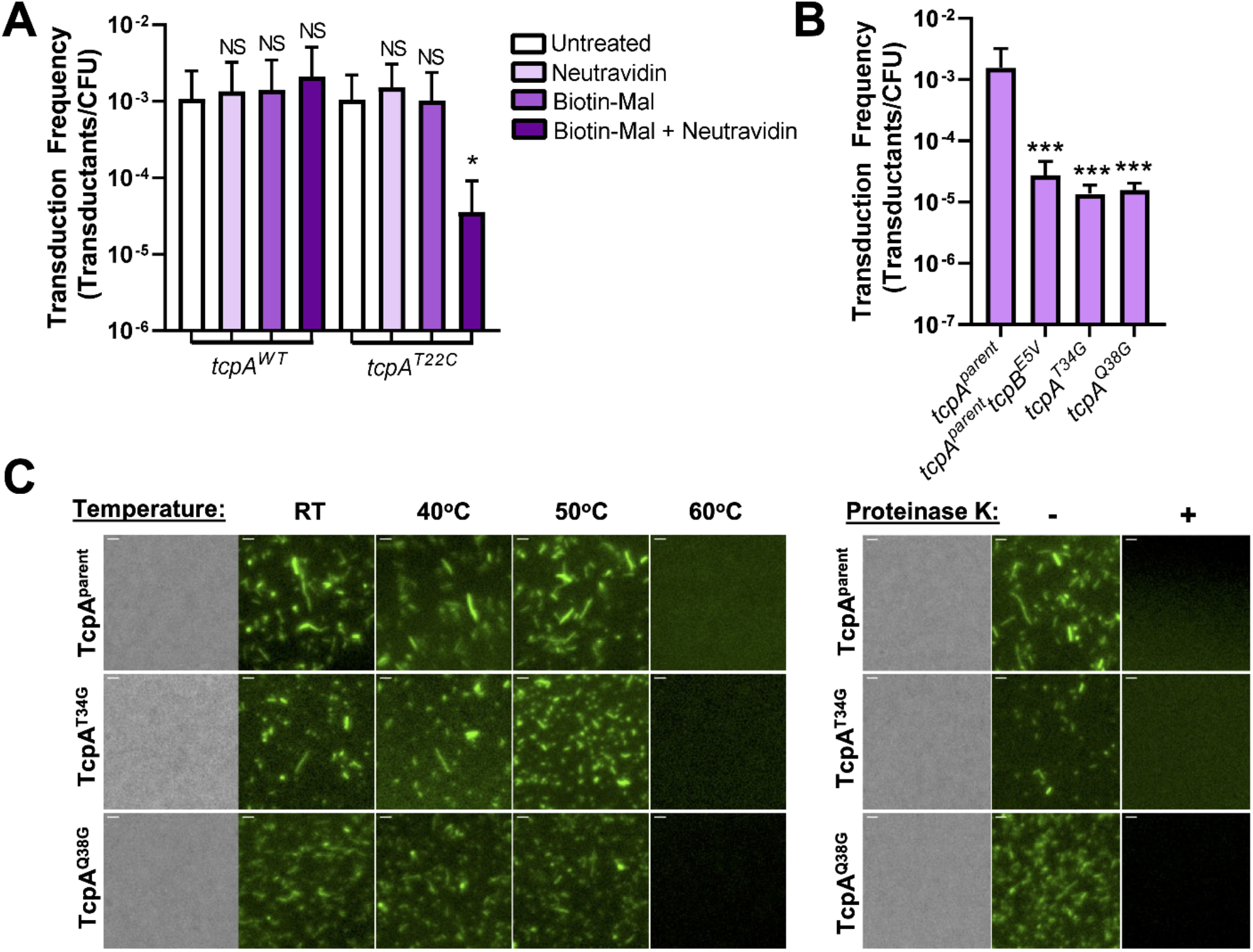
The structure of the pilus fiber is important for motor-independent retraction of the *V. cholerae* toxin co-regulated pilus. **(A-B)** Transduction assays of the indicated strains after the indicated treatment. All strains, *n* = 3. Data are shown as the mean ± SD. Asterisk(s) directly above bars denote comparisons to the parent strain. All comparisons were made by one-way ANOVA with either Dunnett’s post test (**A**) or Tukey’s post test (**B**). LOD, limit of detection; NS, not significant; * = *P* < 0.05, *** = *P* < 0.001. **(C)** Stability assay comparing thermal stability and protease susceptibility of purified AF488-mal labeled pilus filaments from the indicated strains. A representative phase-contrast image is shown to highlight that pilus preps are cell free. Fluorescent images show the presence or absence of intact pili under the indicated conditions. All images are representative of three independent biological replicates. Scale bar, 1 μm.

**Fig. S7.**
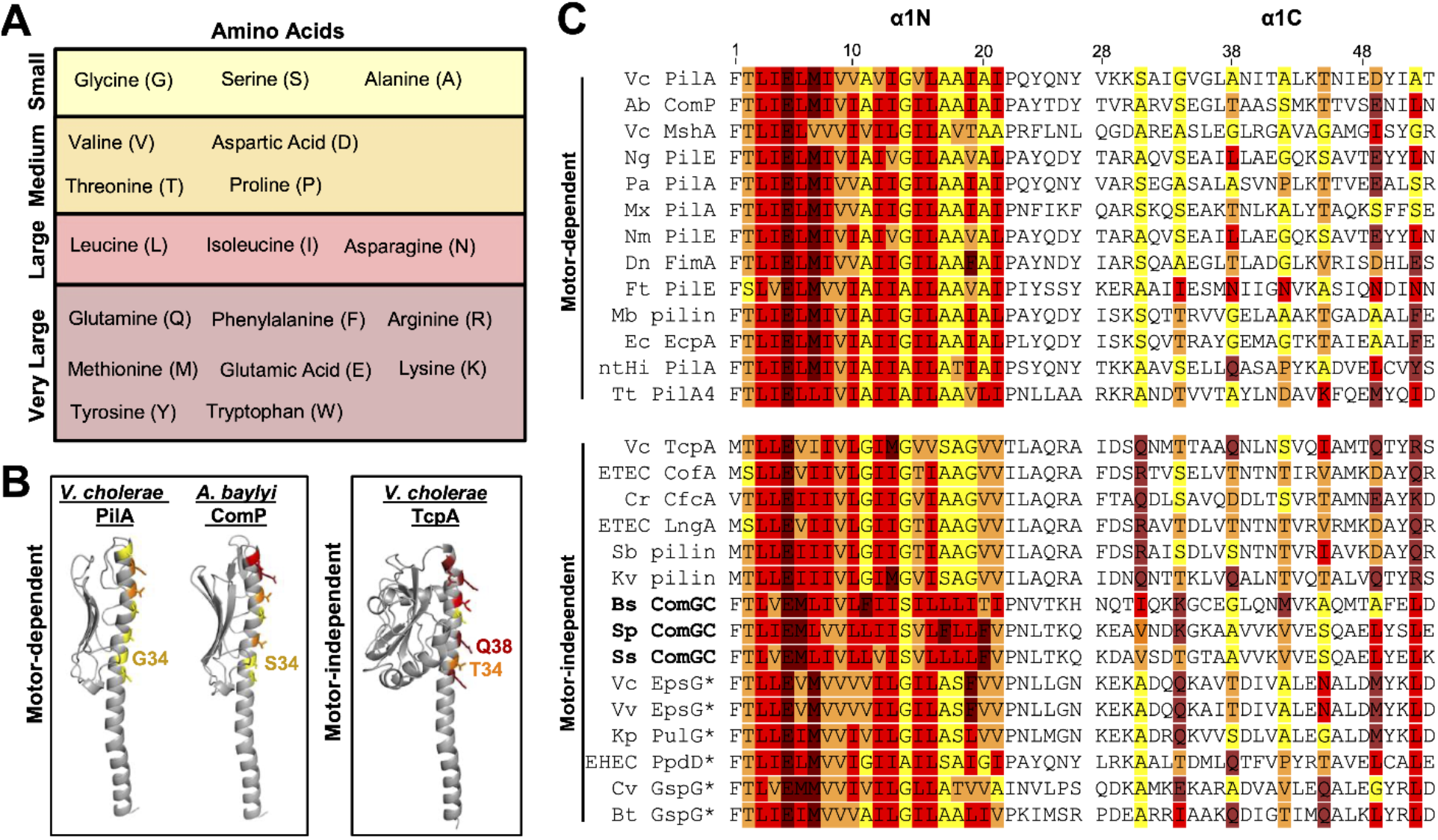
Bulkiness of residues in the α1 helix of major pilins from motor-dependent and motor-independent T4Fs. **(A)** Table of amino acids found in the α-helix of major pilins grouped by size (based on R-group length). **(B)** Structural models of the indicated major pilins showing the size of core-oriented residues in α1C (color coded based on **A**). The positions mutated in this study are demarcated. Phyre2 predicted protein structures were based on threading the major pilin sequence onto the following solved structures: *Vibrio cholerae* PilA (Protein Data Bank (PDB) accession number 1OQW, alignment coverage 80%, confidence 99.9%, identity 38%); *Acinetobacter baylyi* ComP (PDB 1OQW, align. cov. 94%, conf. 99.9%, iden. 32%); *V. cholerae* TcpA, (PDB 1OQV, align. cov. 75%, conf. 99.8%, iden. 77%). **(C)** Sequence alignment of the α1 helix (numbering based on the mature pilin protein) of major pilins from retraction motor-dependent and retraction motor-independent T4Fs. Color coded residues in this region are based on size. The colored residues in the α1N segment (residues 1-27) denote the portion of the pilin that likely interacts with the α1C of neighboring pilin subunits (residues 2-22). The colored residues in the α1C segment (residues 28-53) signify core-oriented amino acids. Pilin sequences in **bold** are Gram-positive Com pili and pilin sequences marked with * are from T2SSs. The GenBank Accession numbers for the protein sequences used are as follows: Vc, *Vibrio cholerae* PilA (NP_232053); Ab, *Acinetobacter baylyi* ComP (WP_004923779); Vc, *V. cholerae* MshA (AAF93582); Ng, *Neisseria gonorrhoeae* PilE (AAA53145); Pa, *Pseudomonas aeruginosa* PilA (WP_058135760); Mx, *Myxococcus xanthus* strain PilA (WP_011555734); Nm, *N. meningitidis* PilE (CAA30557.1); Dn, *Dichelobacter nodosus* FimA (X52403); Ft, *Francisella tularensis* PilE (CAG45522.1);Mb, *Moraxella bovis* pilin (AAA53087.1); Ec, *Eikenella corrodens* EcpA (P35645.1); ntHi, nontypeable *Haemophilus influenzae* PilA (AAX87353); Tt, *Thermus thermophilus* PilA4 (Q72JC0.1); Vc, *V. cholerae* TcpA (WP_001176374); ETEC, enterotoxigenic *Escherichia coli* CofA (WP_110409206); Cr, *Citrobacter rodentium* CfcA (AAO17805); ETEC, enterotoxigenic *E. coli* LngA (AAC33154); Sb, *Salmonella bongori* pilin (WP_138993389.1); Kv, *Klebsiella variicola* pilin (WP_177342149); Bs, Bacillus subtilis ComGC (AFQ58422.1); Sp, *Streptococcus pneumoniae* ComGC (WP_054380926); Ss, *S. sanguinis* ComGC (WP_072073266.1); Vc, *V. cholerae* EpsG (WP_000738789.1); Vv, *V. vulnificus* EpsG (WP_011078941.1); Kp, *Klebsiella pneumoniae* PulG (ABR75617.1); EHEC, Enterohemorrhagic *E. coli* PpdD (WP_025757453.1); Cv, *Caulobacter vibrioides* GspG pilin (WP_058347990.1); Bt, *Burkholderia thailandensis* GspG (EIP89370.1).

---

**Movie S1. There is very little dynamic pilus activity when *pilB* is constantly induced in a Δ*pilTU* strain.**

Pilus dynamic activity was captured by epifluorescence time-lapse microscopy of AF488-mal labeled cells. This time-lapse shows a P_*BAD*_-*pilB^WT^* Δ*pilB pilA^S56C^* Δ*pilTU* strain with constant induction of *pilB^WT^* through addition of arabinose during growth and while imaging. Data are representative of three independent biological replicates. Scale bar, 1 μm.

**Movie S2. Dynamic pilus activity is enhanced when *pilB* is induced in a Δ*pilTU* strain during imaging.**

Pilus dynamic activity was captured by epifluorescence time-lapse microscopy of AF488-mal labeled cells. This time-lapse shows a P_*BAD*_-*pilB^WT^* Δ*pilB pilA^S56C^* Δ*pilTU* strain that was grown without arabinose and *pilB^WT^* expression was induced immediately before imaging. Data are representative of three independent biological replicates. White arrows indicate pilus retraction events. Scale bar, 1 μm.

**Movie S3. Dynamic pilus activity is enhanced when*pilB^M391C^* is induced in a Δ*pilTU* strain during imaging.**

Pilus dynamic activity was captured by epifluorescence time-lapse microscopy of AF488-mal labeled cells. This time-lapse shows a P_*BAD*_-*pilB^M391C^* Δ*pilB pilA^S56C^* Δ*pilTU* strain that was grown without arabinose and *pilB^M391C^* expression was induced immediately before imaging. Data are representative of three independent biological replicates. White arrows indicate pilus retraction events. Scale bar, 1 μm.

**Table S1.** List of important residue positions within the indicated major pilins based on the pre-pilin or mature pilin protein sequence.

| Organism, Major pilin | Pre-pilin position | Mature pilin position |
| --- | --- | --- |
| Vc PilA | G59R | G34R |
| Vc PilA | S67C | S56C |
| Vc PilA | V99A | V74A |
| Vc PilA | G103S | G78S |
| Vc PilA | G105S | G80S |
| Ab ComP | S40R | S34R |
| Ab ComP | T129C | T123C |
| Vc TcpA | T47C | T22C |
| Vc TcpA | T59G | T34G |
| Vc TcpA | Q63G | Q38G |

\*Prepilin positions are relative to the start codon. Mature pilin positions are relative to the first amino acid in the full-length processed pilin protein. Vc, *Vibrio cholerae*; Ab, *Acinetobacter baylyi*.

**Table S2.**
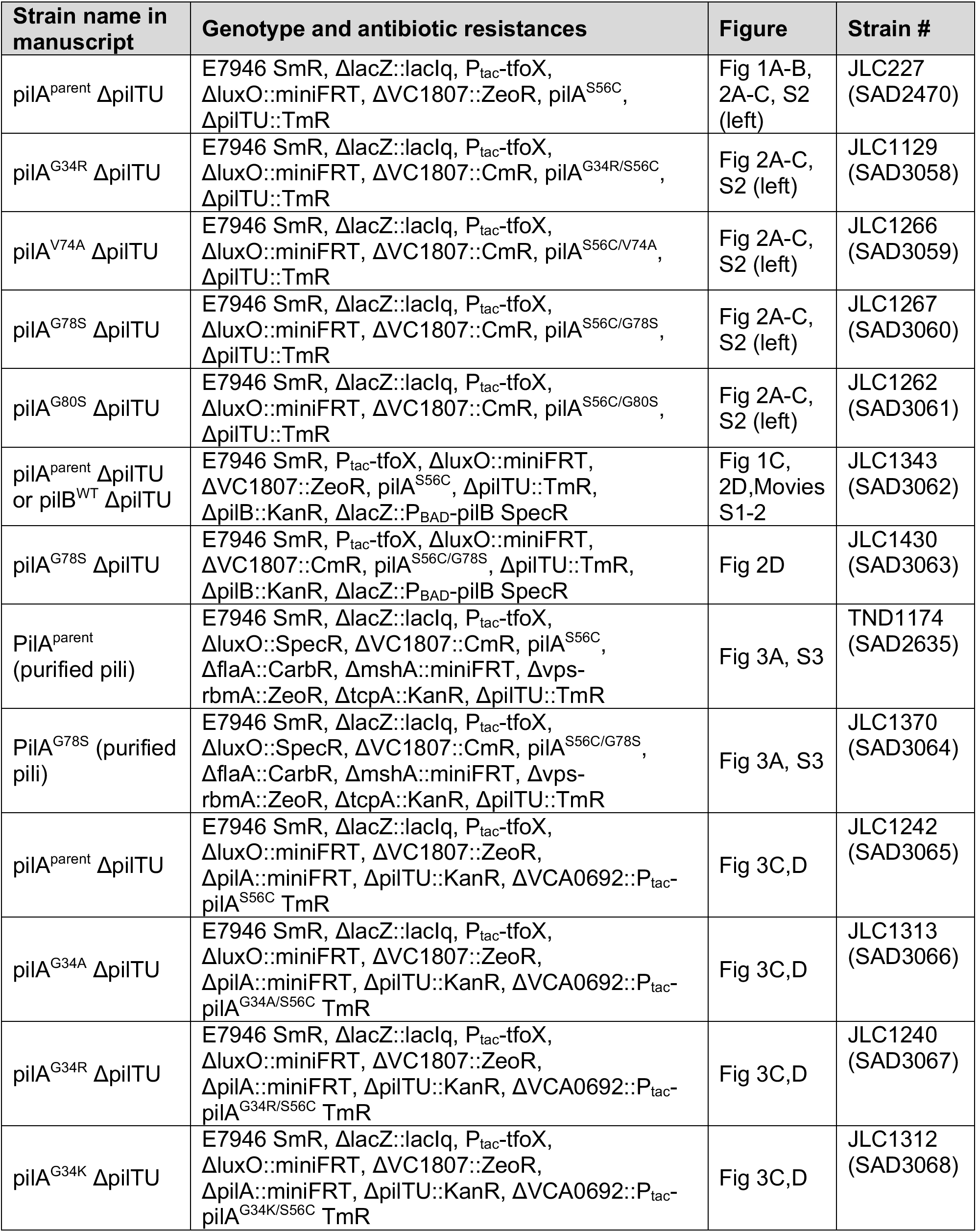
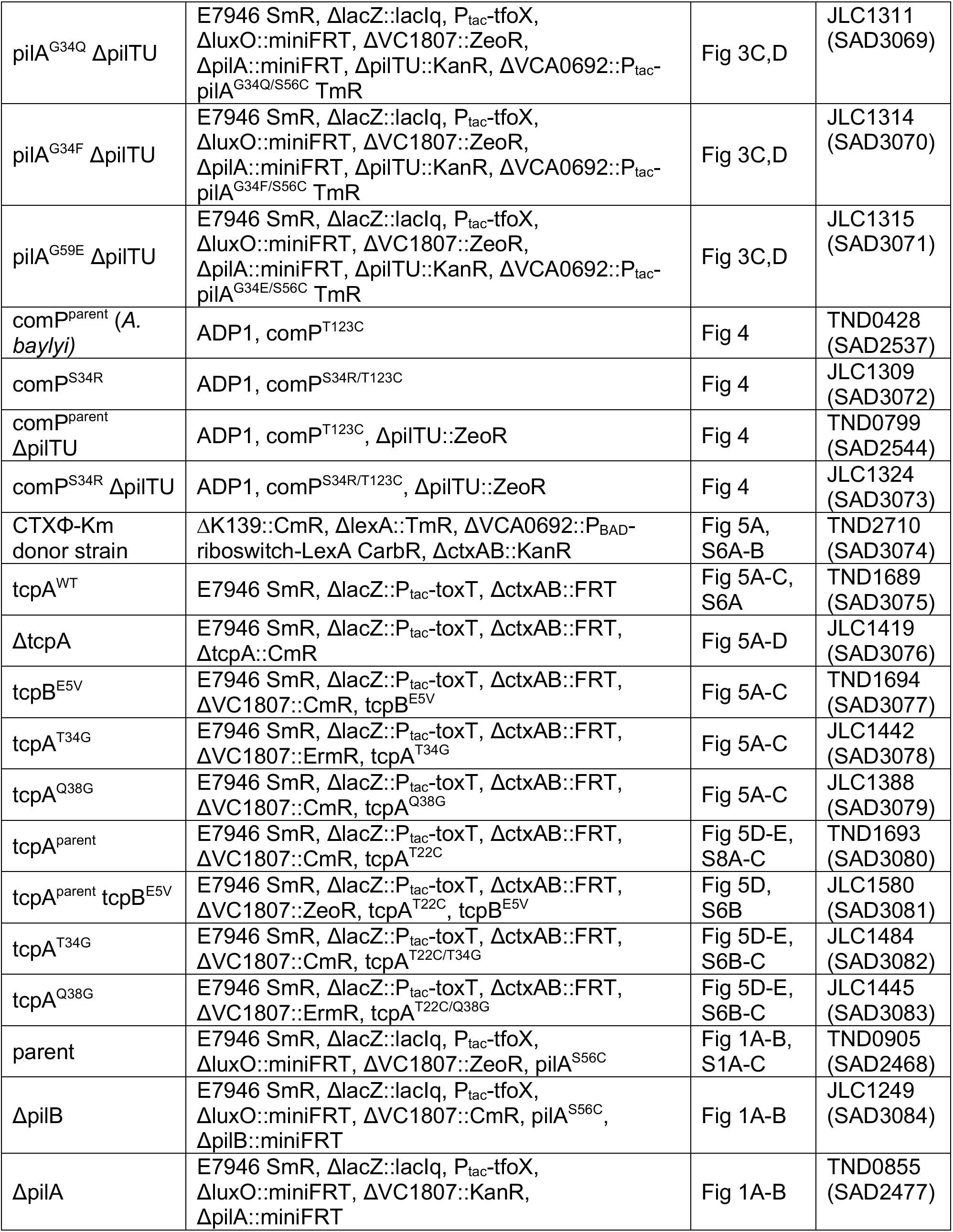
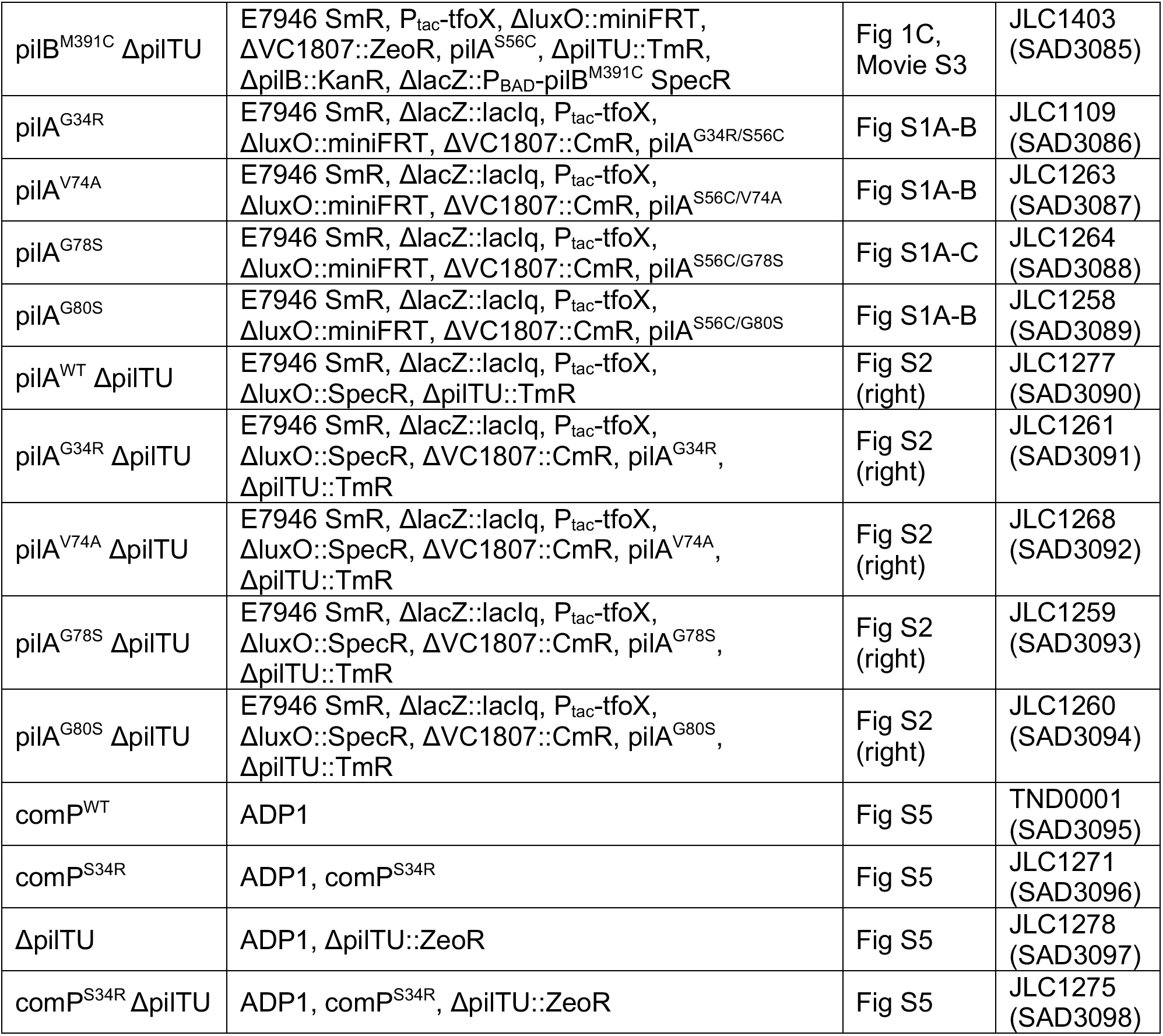
Strains used in this study.

**Table S3.**
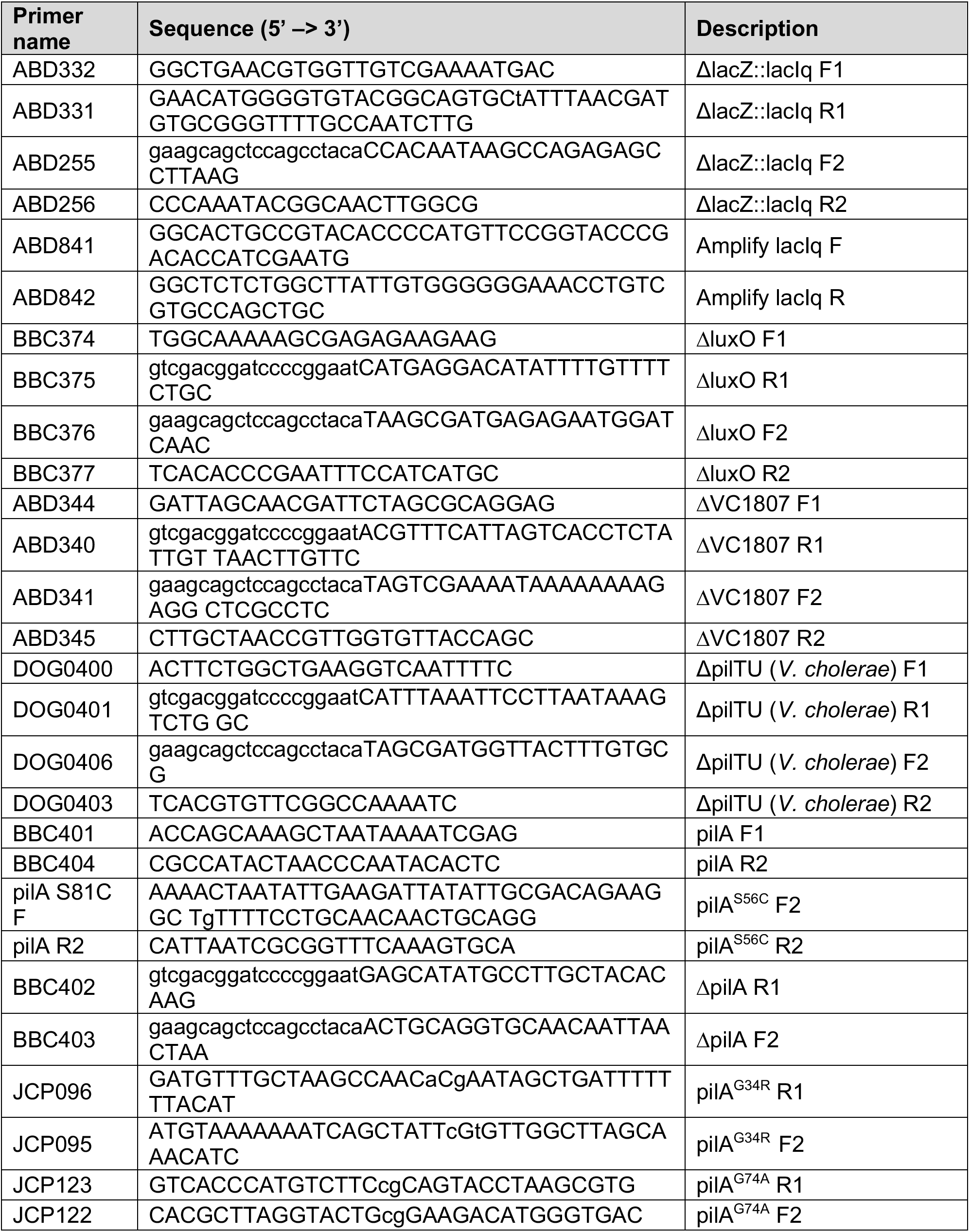
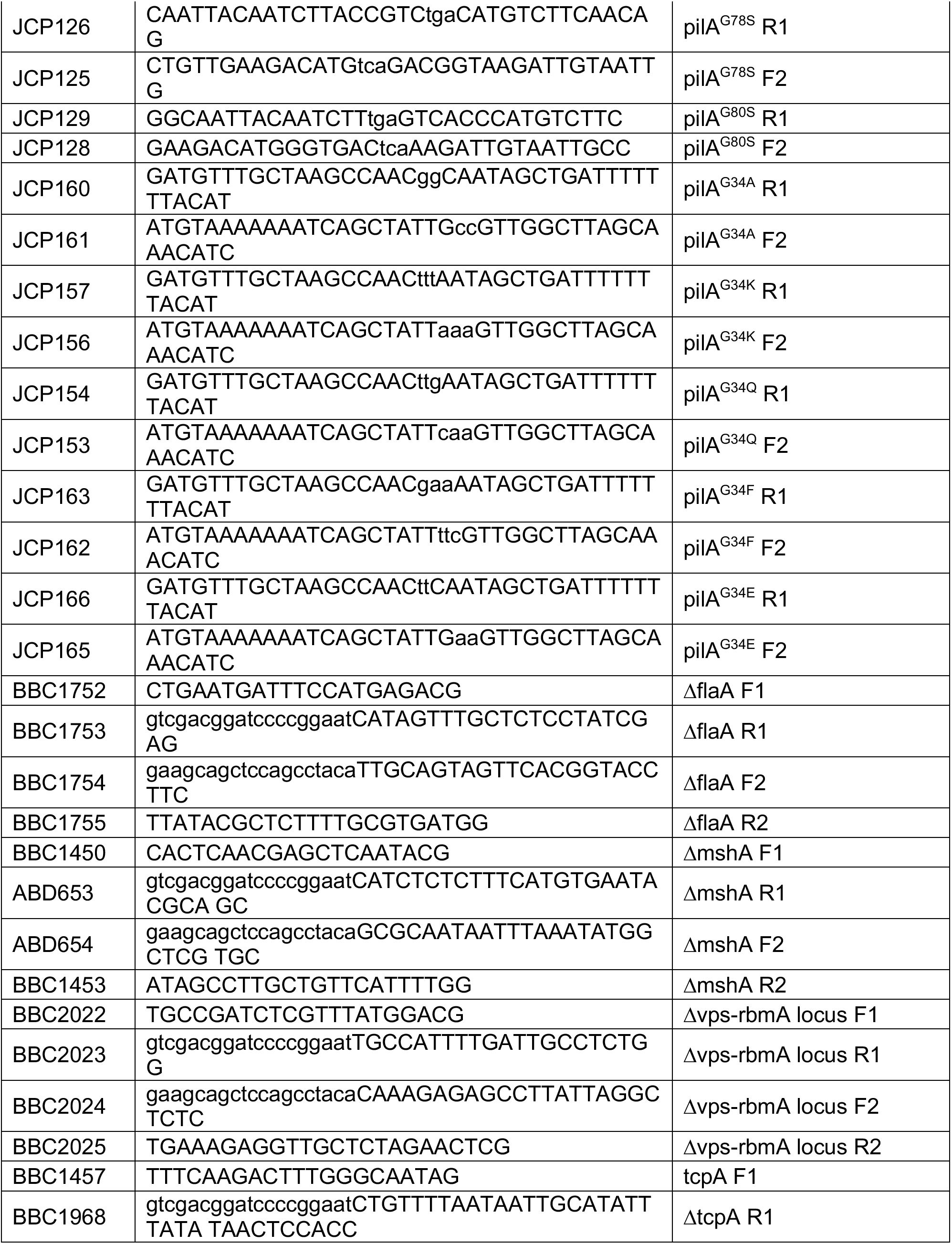
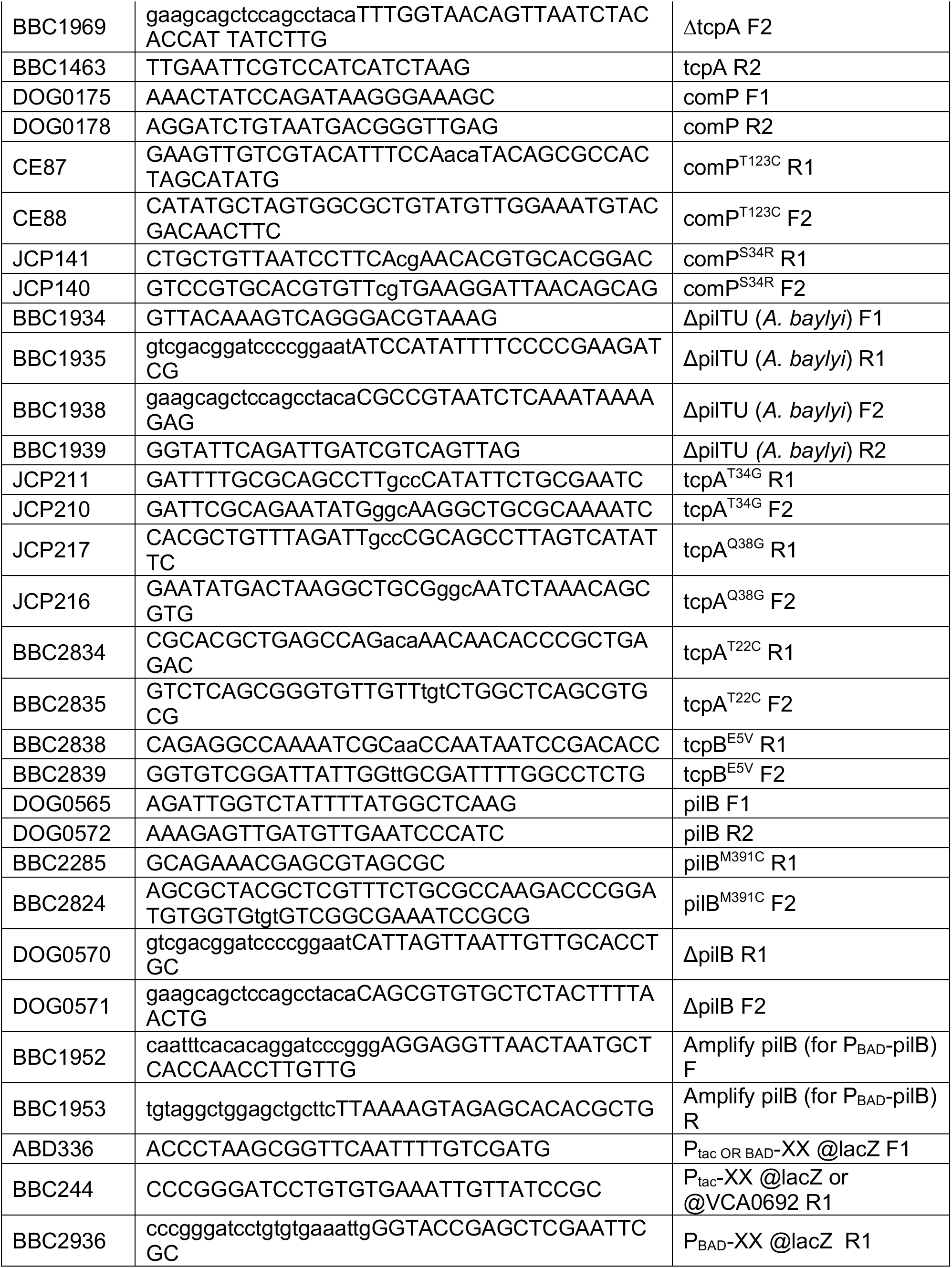
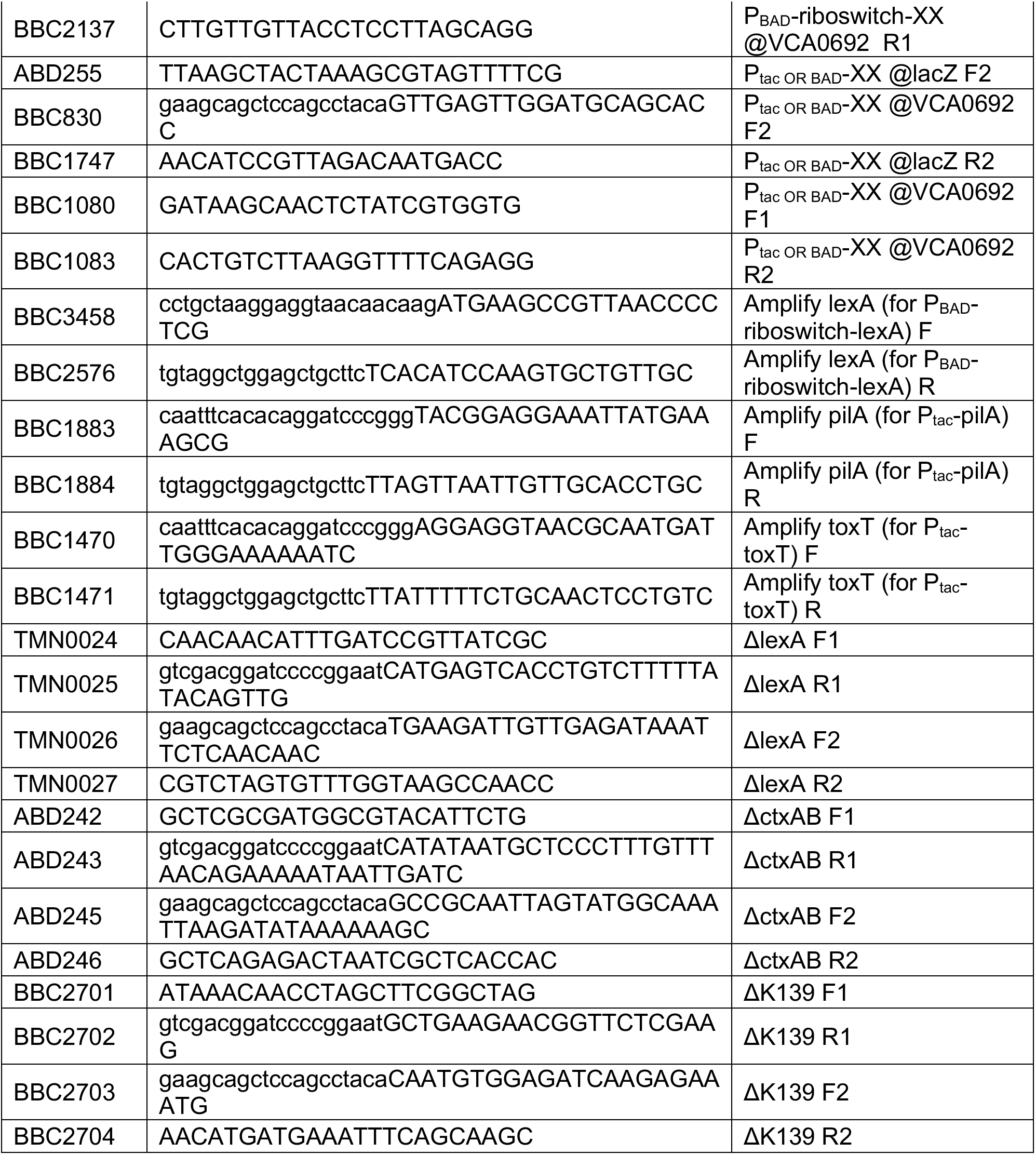
Primers used in this study.

**Table S4.**
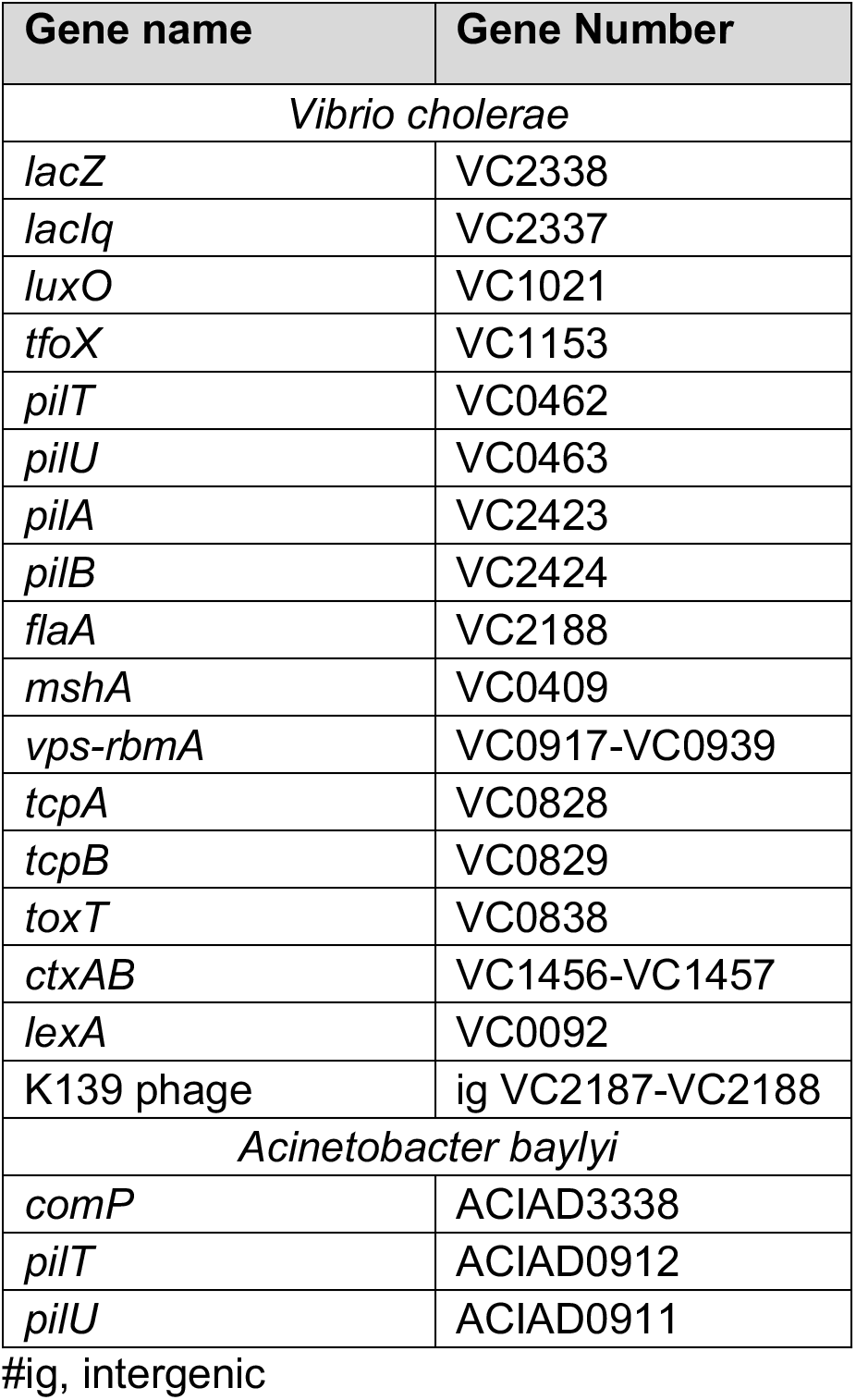
Gene numbers.

**Table S5.**
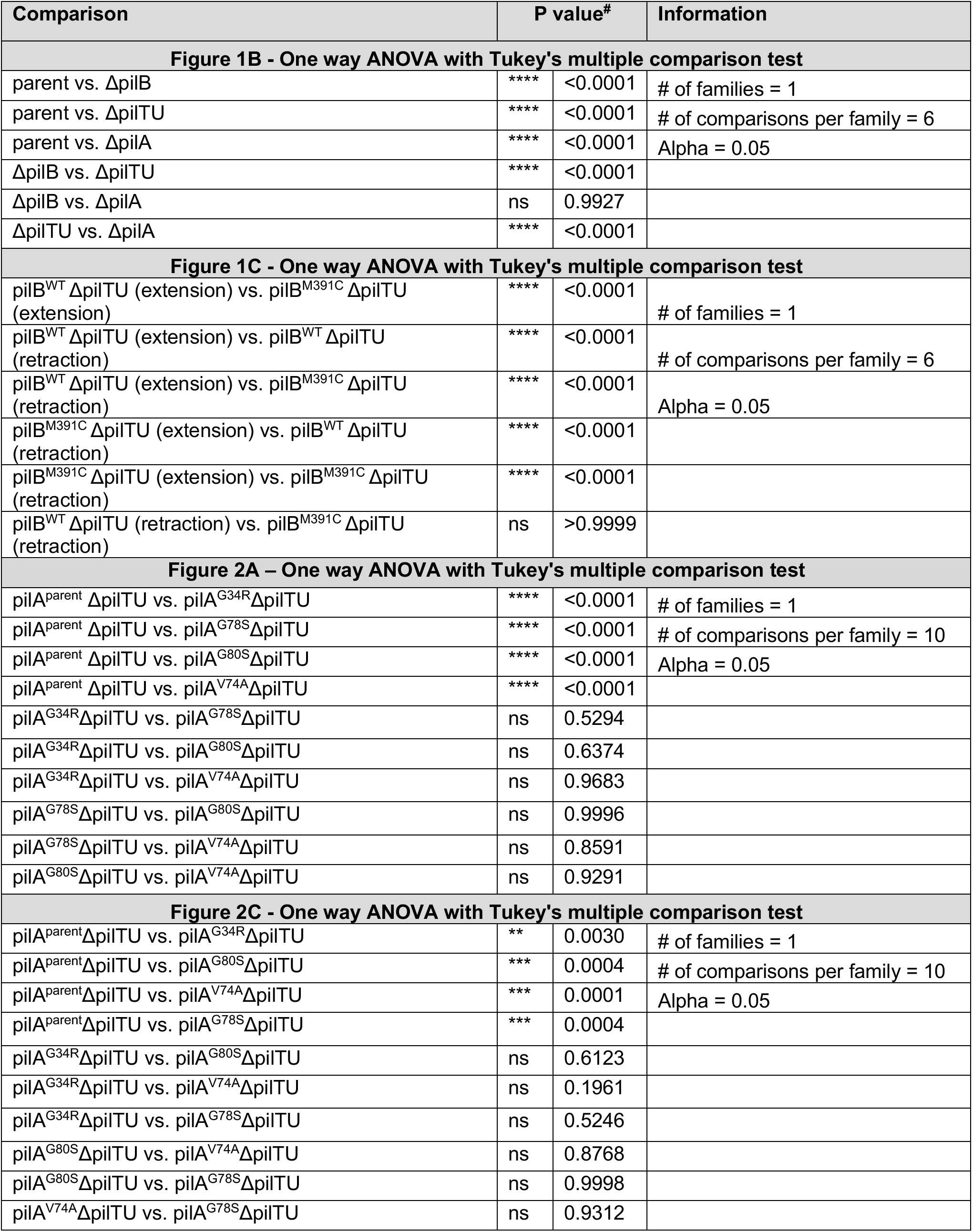
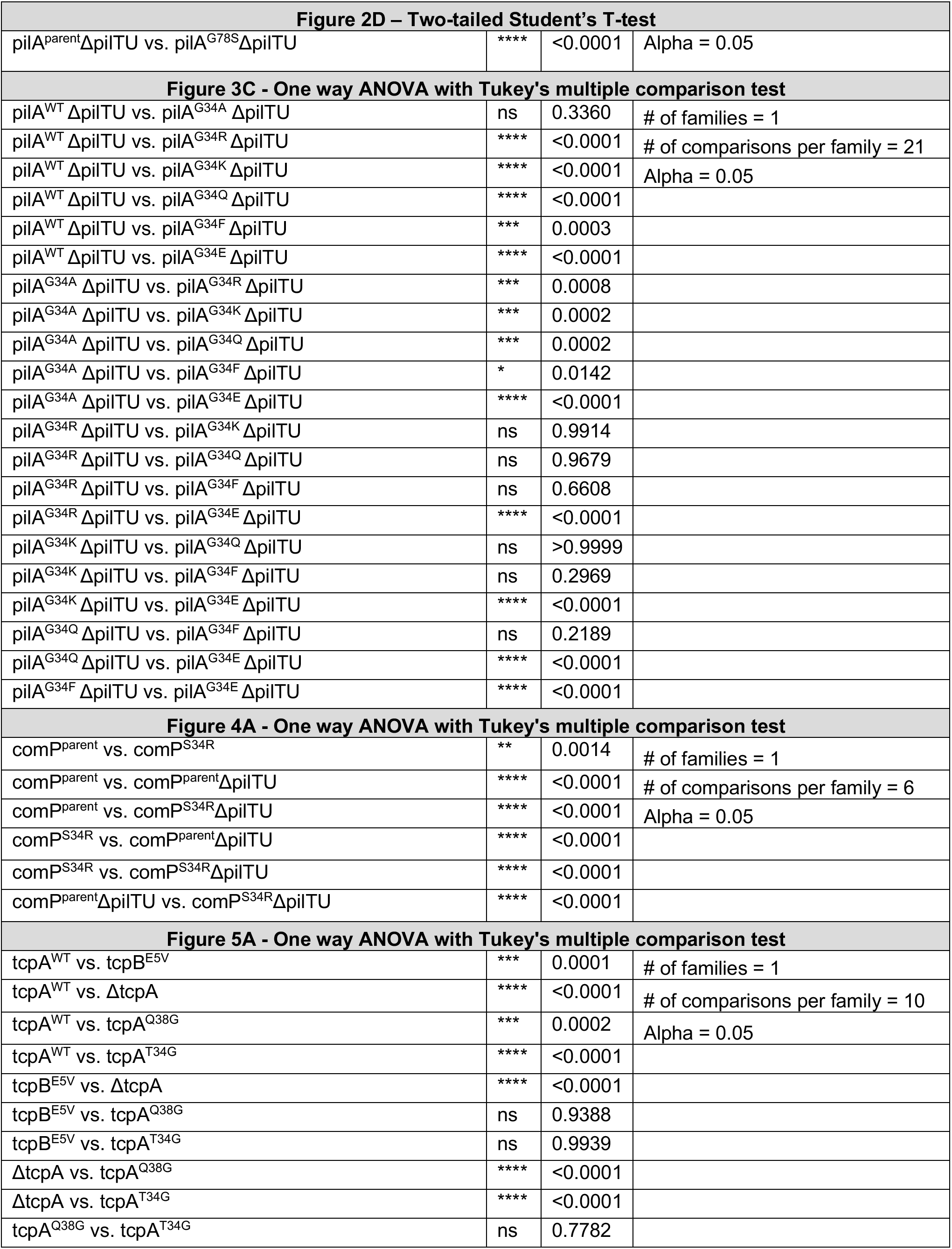
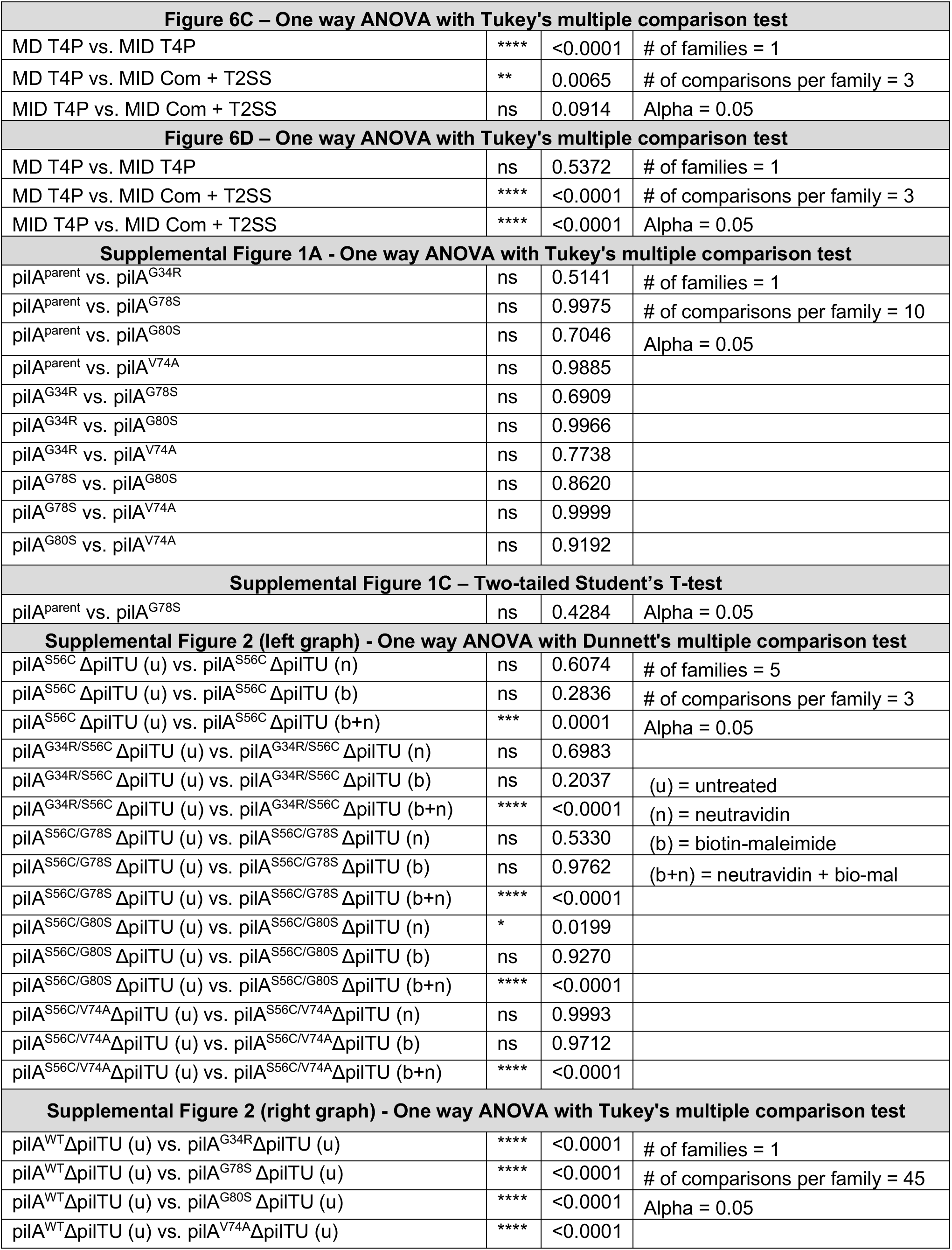
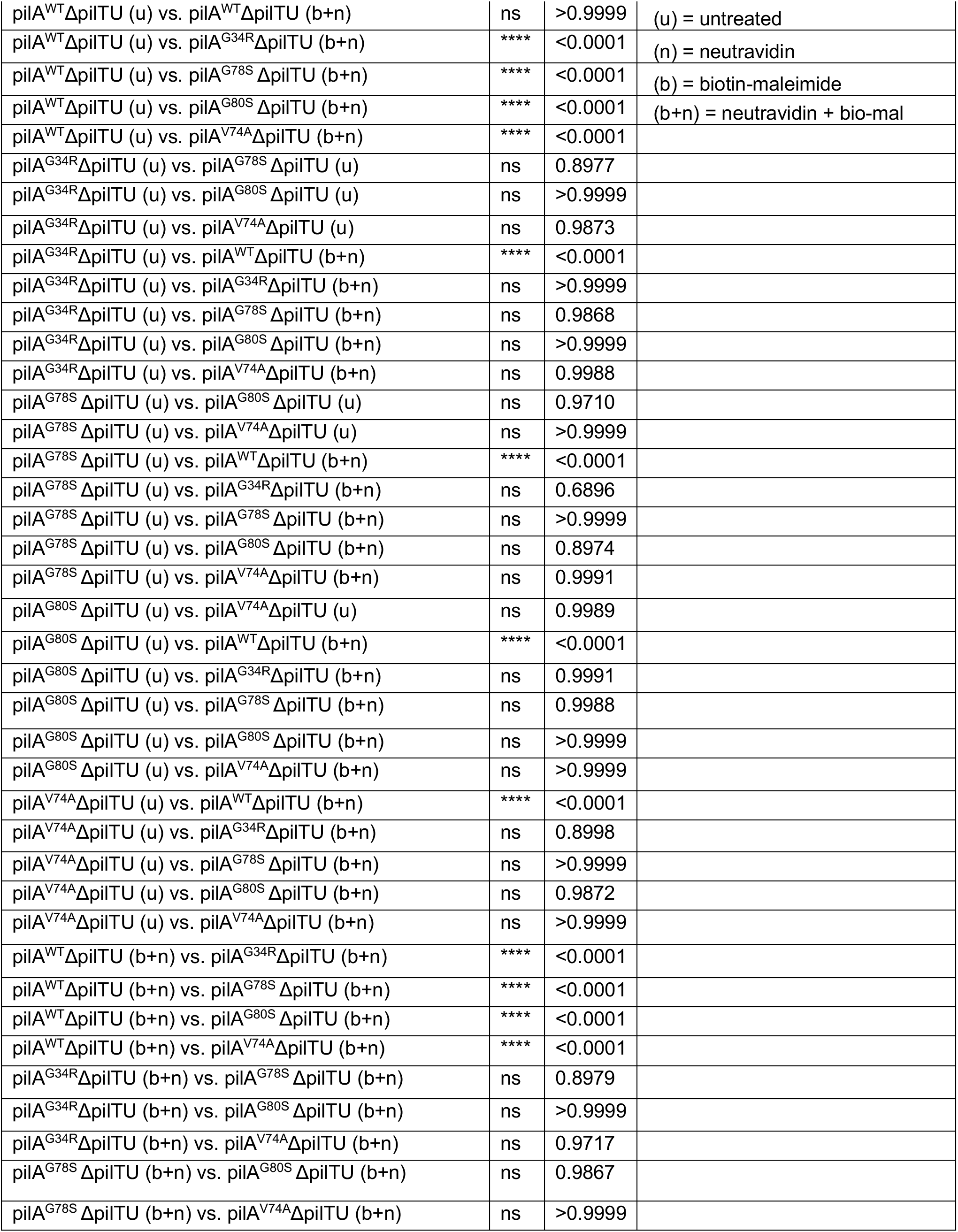
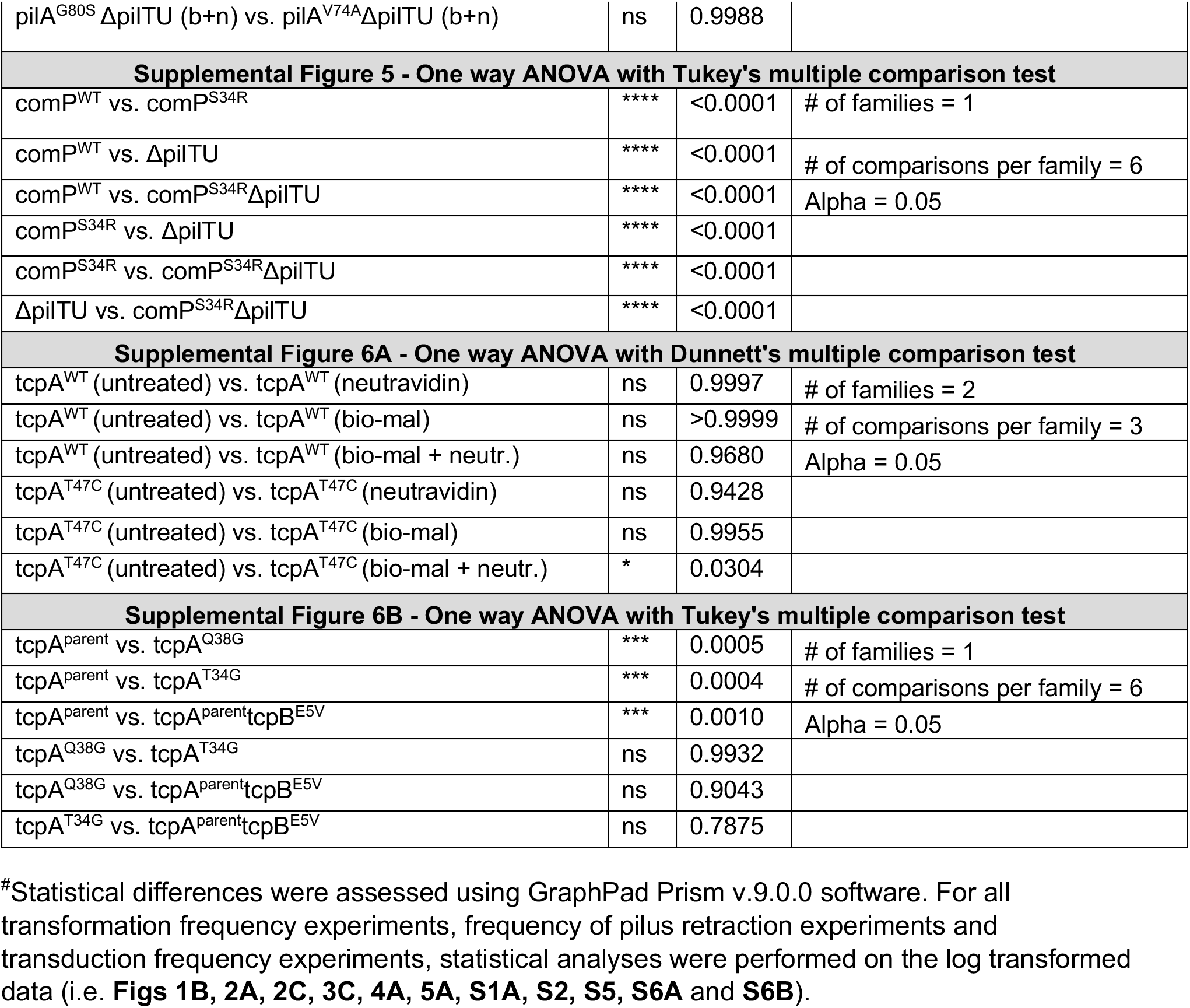
Statistical Comparisons.

